# Targeted and Untargeted Amine Metabolite Quantitation in Single Cells with Isobaric Multiplexing

**DOI:** 10.1101/2024.06.18.599468

**Authors:** Juho Heininen, Parisa Movahedi, Tapio Kotiaho, Risto Kostiainen, Tapio Pahikkala, Jaakko Teppo

## Abstract

We developed a single cell amine analysis approach utilizing isobarically multiplexed samples of 6 individual cells along with analyte abundant carrier. This methodology was applied for absolute quantitation of amino acids and untargeted relative quantitation of amines in a total of 108 individual cells using nanoflow LC with high-resolution mass spectrometry. Together with individually determined cell sizes, this provides the first absolute quantification of intracellular metabolites within individual cells. The targeted method was partially validated for 10 amino acids with limits of detection in low attomoles, linear calibration range covering analyte amounts typically from 30 amol to 120 fmol, and correlation coefficients (R) above 0.99. Using the cell sizes determined during dispensing, millimolar intracellular concentrations were determined. The untargeted approach yielded 249 features that were detected in at least 25% of the single cells, providing modest cell type separation on principal component analysis. Using Greedy forward selection with regularized least squares, a sub-selection of 100 features explaining most of the difference, was determined. These features were annotated using MS2 from analyte standards and accurate mass with library search. The approach provides accessible, sensitive, and high-throughput method with the potential to be expanded also to other forms of ultrasensitive analysis.

## Introduction

As the foundational units of biology, cells exhibit profound heterogeneity, even among seemingly similar cells. This can be caused by environmental perturbations, stochastic variations, differential gene regulation, and mutations. Such intrinsic heterogeneity can lead to diverse biomolecule concentrations, cellular phenotypes, and responses, even in seemingly similar and clonal cells.^1–4^ Studying cells individually sheds light on this cellular heterogeneity and unique attributes, as opposed to relying on bulk averages. This approach is crucial for tackling complex and unresolved problems and advancing our understanding of cellular heterogeneity, development, dynamics, lineages, cell-to-cell interactions, and disease mechanisms. For example, the presence of drug-resistant phenotypes may explain the incomplete eradication of cancer cells by drug treatments. Similarly to the advances in single cell genomics and transcriptomics^5^, progress in single cell metabolomics methodology can provide ways for in-depth studies of cancers like high-grade serous ovarian adenocarcinoma (HGSOC), a common and particularly lethal type of ovarian cancer, as well as the elusive mechanisms driving chemoresistance, their diversity, and distribution between cells.^6^

Despite substantial incentives for single-cell analysis and the availability of sensitive mass spectrometry approaches, typical low picoliter volumes of cells, combined with even the abundant micromolar concentrations of amino acids, results in sub-femtomole quantities of analytes, providing considerable challenge to sample preparation and measurement. Unlike in bulk, single cell analytes cannot be readily enriched (e.g., through Solid Phase Extraction), and unlike DNA or RNA, metabolite amounts cannot be amplified (e.g., using PCR). Combined with sample losses, such as analyte adsorption in above cellular amounts^7^ with common sample preparation approaches, analyte amounts in single cells are orders of magnitude below conventional liquid chromatography (LC) — mass spectrometry (MS) limits of detection (LODs). Improvements in sample preparation methods and LC-MS instrumentation have improved the sensitivity closer towards single cell analysis. For example, unlabeled amino acids have typically been detected with LC-MS from 10 fmol amounts^8,9^, whereas derivatized amino acids (with or without ion pairing compounds) have been reported to have LODs in the low fmol range, owing to their improved ionization efficiency and chromatographic separation.^10–12^ The high sensitivity and selectivity of LC-MS makes it highly competitive compared to non-mass spectrometric methods, such as fluorescence or colorimetric detection of labeled amino acids, which typically yield LODs ranging from low pmol to high fmol for the former, and pmol-scale for the latter.^13^

Direct and hyphenated MS techniques have been applied to single cell analysis thus far. In direct MS methods, metabolites of individual cells have been successfully measured by manual manipulation of single cells into microneedles^14–18^ or by surface desorption methods such as SIMS^19^, DESI^20^, nanoDESI^21^, LAESI^22^, LAAPPI^23^, or MALDI^24^. Even though direct MS methods are fast, their selectivity is heavily dependent on specialized equipment and limited, for example, in the separation of isobaric compounds and in ways to combat ionization suppression. Analyte separation in single cell analysis has thus far been achieved with capillary electrophoresis (CE)^25,26^ for large cells. Nevertheless, the single cell metabolome has remained mostly untapped by scientific research because easily accessible methods having high throughput and providing specific and absolute metabolite quantitation are lacking. The huge chemical space covered by the metabolome, challenging feature annotation, and the requirement of highly specialized instrumentation have deterred both targeted and untargeted single cell metabolomics.

Lately, experiment designs and sample preparation methods towards single cell mass spectrometric analysis have emerged, primarily driven by advancements in single cell proteomics. Notable developments include the introduction of a carrier channel along with multiplexed samples in the BASIL^27^ and iBASIL^28^ methods, as well as SCOPE-MS^29^ and its successor SCOPE2.^30^ Progress has also been made with automating and miniaturizing sample preparation to a few microliters using well plates (mPOP)^31^ or even further to sub microliter volumes on glass slides (nanoPOTS^32^ and nPOP).^33^ Concurrent advancements have been achieved also with scaling down the liquid phase separation as well as electrospray ionization to nanoscale, providing higher ionization efficiency and improved sensitivity.^34–38^

In this work, we developed a sensitive quantitation method using isobaric multiplexing with a carrier that is suitable for single cell analysis. Using absolute quantitation approach with cell size determination, this provides the first intracellular concentration determination of amino acids in single cells. We further show the applicability of the method to untargeted relative quantification of metabolites. We measured amine metabolites in individual cells of the high-grade serous ovarian adenocarcinoma (HGSOC) cell line TYK-nu and its cisplatin-resistant variant TYK-nu.CP-r, comparing them to each other and to human embryonic kidney 293 (HEK-293) cells. Sufficient sensitivity was achieved using a carrier analyte, miniaturized sample preparation, and optimized data acquisition with nLC-MS. For cell sorting, we employed the CellenONE platform, which provides a microscope image and cell size measurement, required to determine intracellular analyte concentrations, for each sorted cell. Sample preparation was carried out with a pipetting robot capable of working with low microliter volumes, thus ensuring robust, traceable, and repeatable analysis of single cells.

## Results and Discussion

To improve sensitivity for single cell analysis and mitigate sample losses, amine metabolites in single cell samples were labeled using isobaric Tandem Mass Tags (TMT)^39^, a set of isotopically different MS2 resolvable labeling reagents (isotopomers), and multiplexed together with analyte-abundant carrier in a concept similar to that used in single cell proteomics.^27,28,40,41^ Each LC-MS run composed of nine individual samples: six individual single cells, analyte-abundant carrier, blank with no analytes and blank with internal standards (IS-blank) to manage false positives, each labelled with a different TMT isotopomer. Since the labeled analytes in individual samples and the carrier are chemically similar, the sample losses are primarily mitigated by the carrier. Furthermore, their superimposed MS1 peak provides a robust MS1 signal, enabling confident targeted Parallel Reaction Monitoring (tPRM) isolation window construction for targeted metabolite analysis, as well as for untargeted metabolomics employing intensity dependent (e.g. top N) precursor selection in DDA (Data Dependent Acquisition). The individual analytes can be distinguished from isobaric MS1 ions using MS2 analysis, as the isotopomer labels fragment to their unique *m/z* reporter ions. This concept is illustrated in Figure 1.

**Figure 1.**
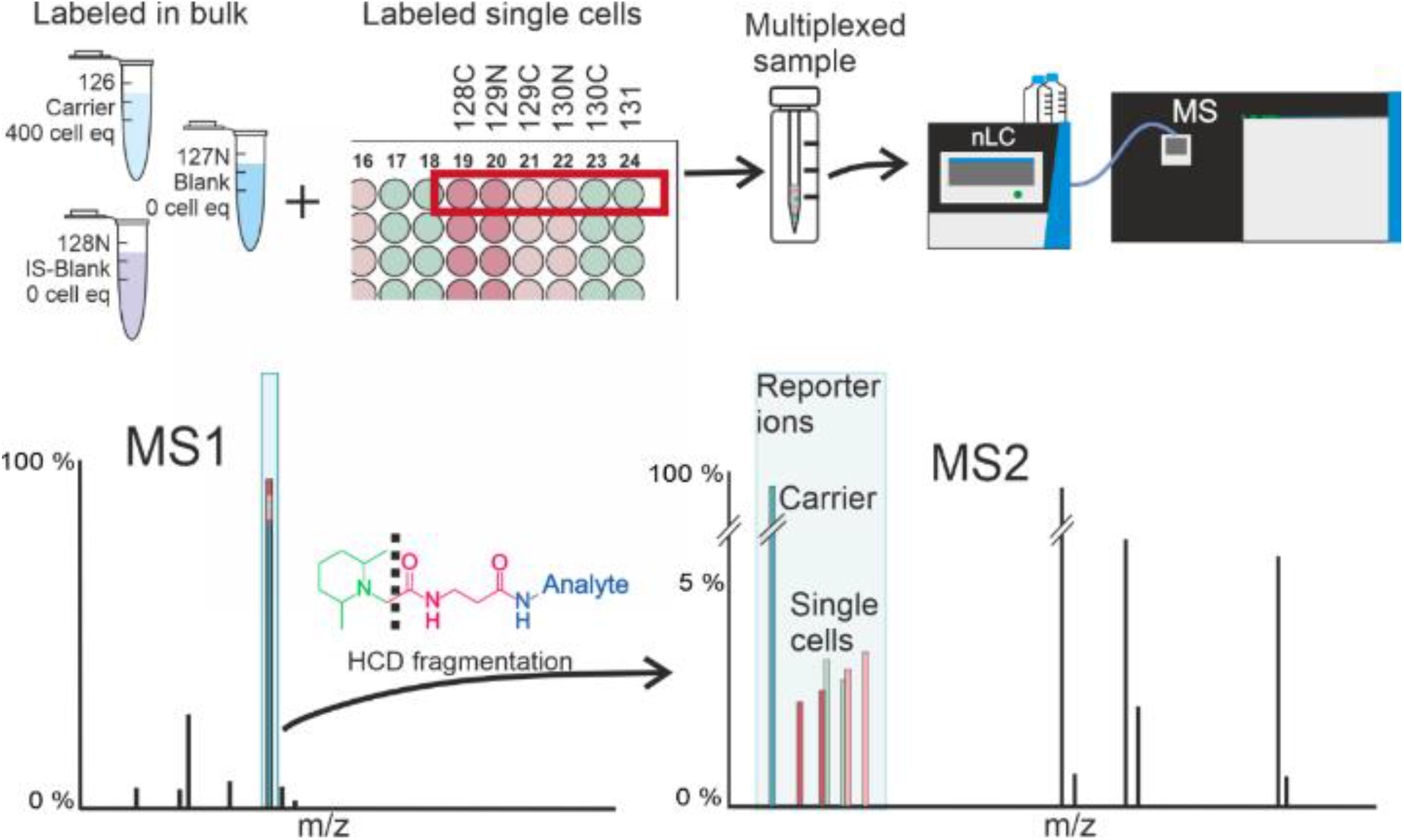
Isobaric labeling concept for single cell samples. Multiplexed sample consists of six individual single cells prepared and labeled individually on a 384-well plate along with analyte abundant carrier, blank, and IS-blank prepared and labeled in bulk. From the ten TMT10plex reagents, nine (126, 127N, 128N, 128C, 129N, 129C, 130N, 130C, and 131) were used, whereas one (127C) was not, due to isotope impurity effect of analyte abundant carrier. These multiplexed samples were measured using nLC-MS2, where each isobarically labeled analyte from the multiplexed sample produces a superimposed MS1 peak that can be decomposed to sample reagent (126-131) specific reporter ions with HCD fragmentation and quantitatively measured with high resolution MS.

### Absolute quantitation method for amino acids in single cells

The TMT-labeled amino acids were observed in MS1 analysis as singly labeled and singly protonated [M+1TMT10+1H]^+1^ ions, except cystine and lysine, which both contain two amine groups, were observed as doubly labeled and doubly protonated [M+2TMT10+2H]^+2^ ions, similarly to previous research.^42,43^ These labeled analyte ions were detected with mass accuracies of below 0.73 mDa. Product ion spectra of the protonated molecules of TMT labeled amino acids were similar to our previous research.^42^ Most prominent product ions were intense TMT10plex isotopomer-specific reporter ions. Other observed product ions were formed by loss of water and carboxylic acid, as well as dissociation of amide bonds. The absolute quantitation method was achieved by measuring TMT10plex isotopomer-specific reporter ions of amino acids and their heavy isotope internal standards with a tPRM approach based on analyte ions detected in LC-MS1 and their retention times.

For chromatography, we employed nLC along with nanoESI for the quantitation of TMT-labeled amino acids in single cells due to its enhanced sensitivity compared to conventional UHPLC-ESI, a benefit derived from both the nanoscale liquid chromatography and the nanoESI source.^44^ From the initial 17 analytes in the amino acid standard, four polar amino acid analytes (serine, arginine, glycine, and histidine), however, exhibited either poor retention time stability or inadequate retention on the reversed phase column, preventing confident quantitation. Although quantifiable, the poor retention of polar analytes can be also observed with aspartic acid in the product ion EICs of TMT reporter ions (m/z 126.1277) of labeled 120 fmol analyte amounts seen in Figure 2. Among the remaining 13 analytes, proline transition presented interference with some TMT10plex reporter ions, preventing quantitation, and the method lacked sufficient sensitivity for low concentrations of alanine and threonine. Consequently, 10 analytes were quantifiable at single-cell concentrations (Table 1). Figure 2 shows the extracted product ion chromatograms of ten TMT10plex labeled amino acids (120 fmol injected) as an example.

**Table 1.** Validation of the LC-MS2 method for the analysis of amino acids in single cells. The calibration curve was weighed using 1/c. Quantification repeatability was measured at 0.3 fmol and 3 fmol sample amounts that roughly correspond to expected 1mM and 0.1 mM amino acid concentrations in a cell with a volume of 3 pL.

| Analyte | Rt (Min) | FWHM (min) | Rt (RSD %) | LOD<sup>[a]</sup> (amol) | LOQ<sup>[a]</sup> (amol) | Calibration range (fmol) | R | Cal points (n) | 0.3 fmol (RSD %) | 3 fmol (RSD %) |
| --- | --- | --- | --- | --- | --- | --- | --- | --- | --- | --- |
| Aspartic acid | 13.40 | 2.15 | 12.6 | 6.9 | 23.1 | 0.03-120 | 0.9976 | 6 | 16.6 | 3.1 |
| Cystine | 19.60 | 0.15 | 0.6 | 9.0 | 30.0 | 0.15-120 | 0.9998 | 6 | 7.9 | 3.8 |
| Glutamic acid | 16.09 | 0.33 | 1.4 | 6.1 | 20.2 | 0.03-120 | 0.9973 | 6 | 15.9 | 7.3 |
| Isoleucine | 24.88 | 0.13 | 0.8 | 2.3 | 7.5 | 0.03-120 | 0.9985 | 6 | 9.3 | 4.0 |
| Leucine | 25.56 | 0.11 | 1.1 | 3.4 | 11.3 | 0.03-120 | 0.9982 | 7 | 55.0 | 2.6 |
| Lysine | 18.39 | 0.09 | 0.5 | 2.6 | 8.6 | 0.03-6 | 0.9917 | 5 | 18.3 | 6.3 |
| Methionine | 21.57 | 0.20 | 0.8 | 6.4 | 21.4 | 0.03-6 | 0.9916 | 6 | 12.9 | 5.9 |
| Phenylalanine | 26.91 | 0.05 | 0.1 | 1.1 | 3.8 | 0.03-120 | 0.9990 | 7 | 14.3 | 6.3 |
| Tyrosine | 21.52 | 0.14 | 0.6 | 2.6 | 8.6 | 0.03-120 | 0.9986 | 7 | 15.6 | 6.4 |
| Valine | 20.96 | 0.18 | 0.7 | 5.1 | 17.1 | 0.03-120 | 0.9994 | 5 | 17.0 | 3.7 |

[a] LOD and LOQ were calculated based on noise on a peak free section of reporter ions within analyte tPRM transitions.

**Figure 2.**
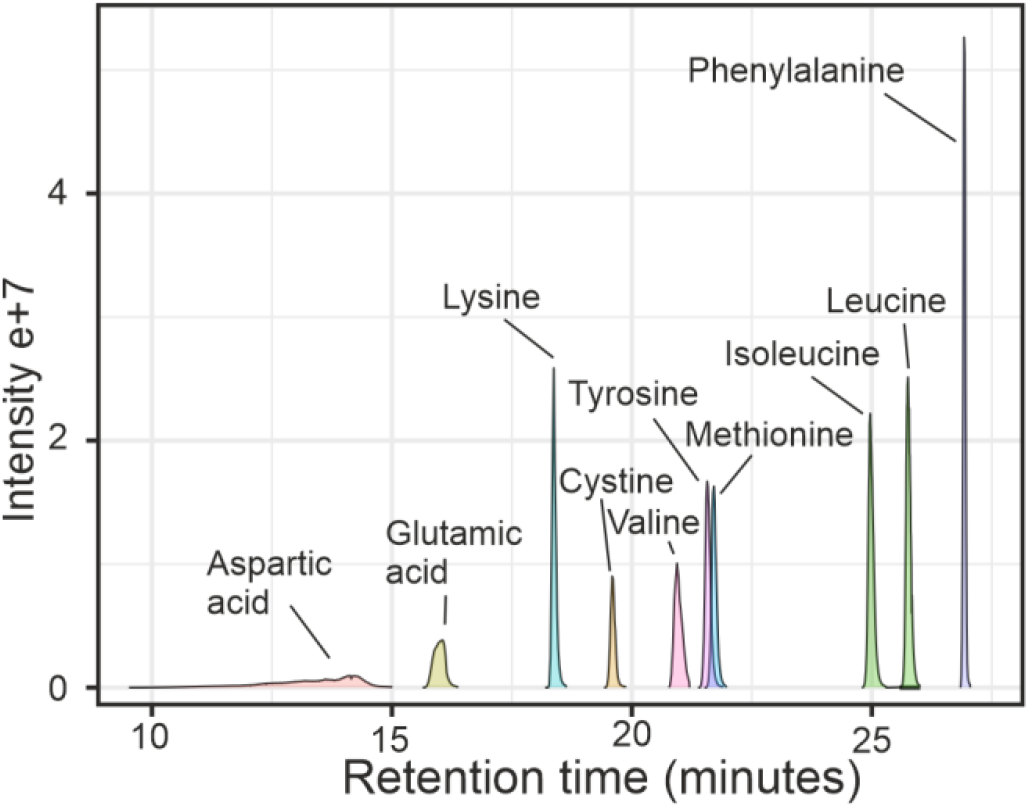
MS2 EICs of integrated reporter ion (m/z 126.1277) peaks of tPRM acquisition of TMT10plex isotopomer labeled 120 fmol analyte amounts from multiplexed calibration sample.

The chromatographic method of the 10 selected TMT-labeled amino acids ensured minimal analyte co-elution, baseline separation of isobaric isoleucine and leucine with peak resolution (Rs) of 1.44, and reproducible retention times with RSDs below 2.5% (excluding the poorly retained but quantifiable labeled aspartic acid with retention time RSD 12.6%). Furthermore, minimal co-elution ensured a sufficient number of scans per chromatographic peak for a quantitative method.

The LOD and LOQ were calculated based on signal-to-noise (S/N) ratios of 3 and 10, respectively, where the noise was estimated from the peak-free section of TMT reporter ion chromatograms of the tPRM transitions. The lowest concentrations in calibration samples were above the highest LOQ value. These LOQs, in the low tens of attomoles range, are suitable for analyzing common amine metabolites in single cells. An average single cell of 3 pL volume with an analyte abundance of 100 µM would require a LOQ below 0.3 fmol, unreachable with common amino acid analysis approaches. For example, unlabeled amino acids have typically been detected from 10 fmol amounts^8,9^, whereas derivatized amino acids (with or without ion pairing compounds) have been reported to have LODs in the low fmol range, owing to their improved ionization efficiency and chromatographic separation.^10–12^ The LODs of amino acids have been improved down to 0.1 fmol by utilizing labels enjoying permanent charge, such as quaternary ammonium compounds, but with tradeoff on the chromatographic performance.^45,46^ Similar LODs have been reached using sample multiplexing with isobaric labeling^47^ without the chromatographic challenges associated with permanent charge reagents.^48^ The measured LOQs in low tens of attomoles here are among the most sensitive LC-MS methods to quantify amino acids. Non-mass spectrometric methods, such as fluorescence or colorimetric detection, of labeled amino acids typically yield LODs ranging from low pmol to high fmol for the former, and pmol-scale for the latter.^13^

Calibration ranges were prepared to cover absolute sample amounts of 0.03-120 fmol (roughly from 10 µM to 40 mM cellular concentrations with 3 pL cell volume) by using seven multiplexed calibration samples. Calibration lines for quantitation were constructed using a 1/x weighted least squares linear regression method. In this approach, the ratios of the analyte peak areas to the internal standard peak area (y) were plotted against the nominal analyte concentrations (x), with the weighting scheme placing greater emphasis on the lower concentration points on the calibration line. Method linearity was assessed from calibration residuals. An example of raw MS2 extracted ion chromatograms (EIC) of TMT10plex labeled cystine reporter ions used for calibration signal and the constructed calibration line is shown in Figure 3, and all calibration lines are in Figure S1.

**Figure 3.**
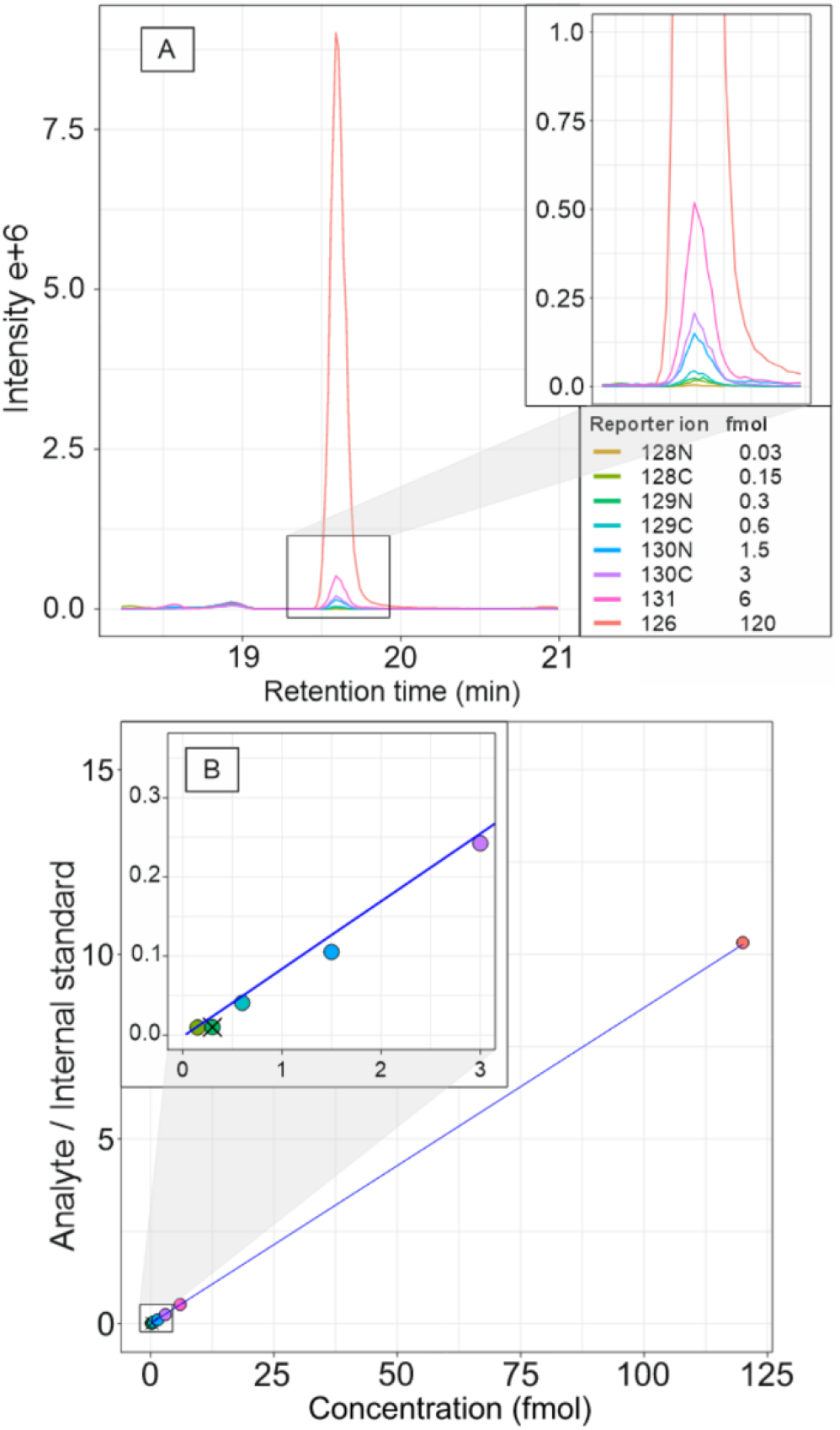
A) EICs of TMT10plex reporter ions of cystine (TMT10plex-cystine precursor, *m/z* 350.1824) from multiplexed calibration sample with amounts of 0.03-120 fmol. B) The constructed calibration line of integrated reporter ion chromatographic peaks after isobaric impurity correction and internal standard normalization. 0.03 fmol sample was removed as being below LOQ and 0.3 fmol that was an outlier with most analytes (Grubb’s outlier test on residuals). These were subsequently excluded from the calibration.

The correlation coefficients of calibration lines were all above 0.99, indicating good linearity within the calibration range for all analytes. Calibration ranges were from 30 amol (0.03 fmol) for all analytes except cystine (60 amol) to the upper limit of quantitation of 120 fmol for all analytes except lysine and methionine (6 fmol), still indicating sufficient analytical range for single cell analysis.

Method repeatability was assessed using seven replicates at 0.3 fmol and 3 fmol analyte amounts. These amounts roughly correspond to typical intracellular amino acid concentrations^49–51^ of 100 µM and 1 mM, respectively, for a mammalian cell with a volume of 3 pL.^52–54^ Examples of the reporter ion chromatograms for lysine (at 0.3 fmol sample amount), are shown in Figure 4.

**Figure 4.**
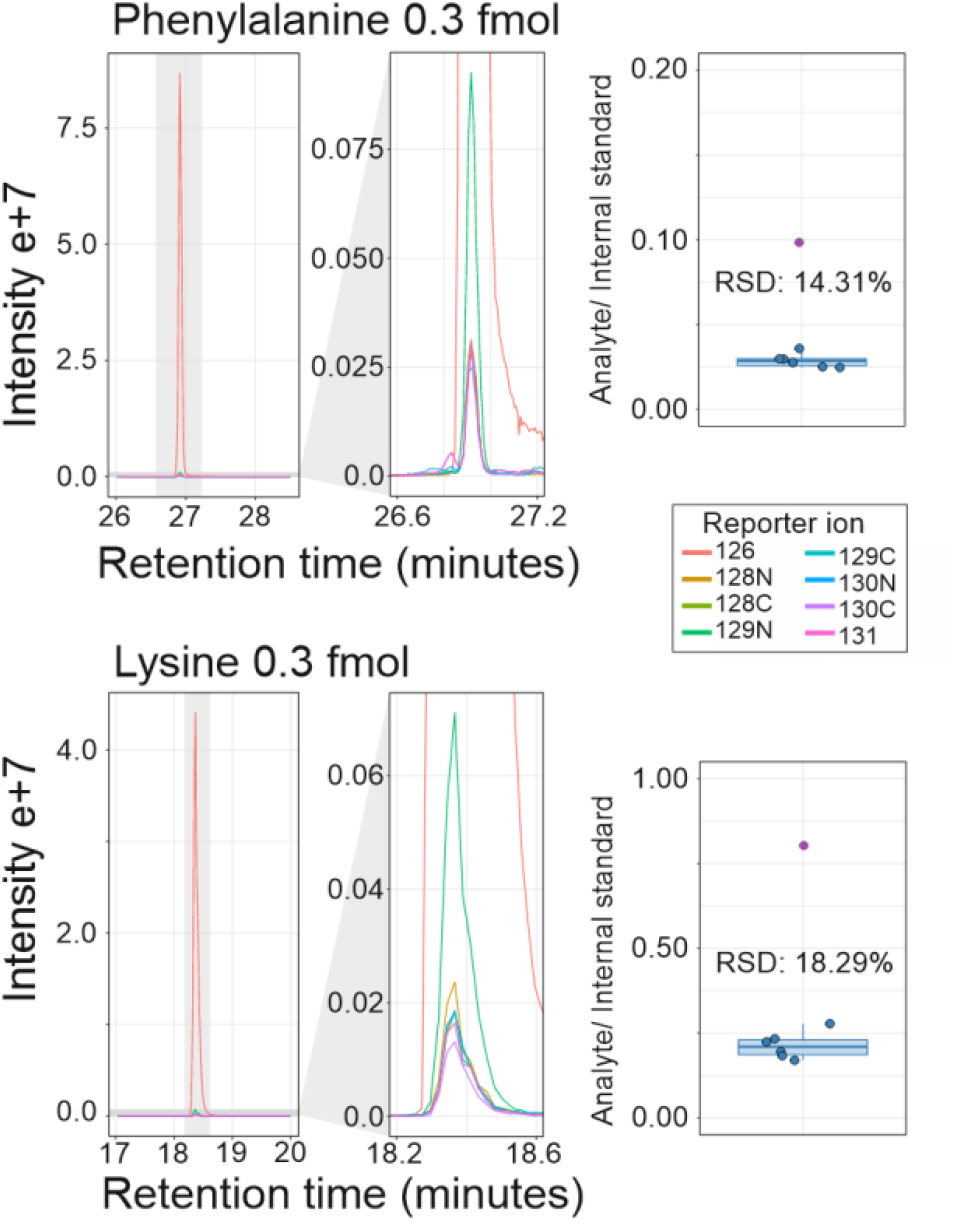
EICs of TMT10plex reporter ions of lysine and phenylalanine replicates at 0.3 fmol sample amount along with the carrier. One sample labeled with 129N (resulting in specific fragment m/z 129.131471) is an outlier. Also individual analyte/IS ratios and resulting relative standard deviation (RSD) are shown excluding the 129N outlier (purple).

Method repeatability RSD with 3 fmol ranged from 2.6% to 7.3% with the average of 4.9%, and with 0.3 fmol from 7.9% to 55% with the average of 18.3% (Table 1), highlighting good method repeatability at cellular sample amounts.

In our work, the high sensitivity required for single cell analysis was achieved through several strategies. We labeled polar analytes with a nonpolar reagent that includes an amine moiety easily ionizable with sensitive nanoflow liquid chromatography-nanoelectrospray ionization. Additionally, we used carrier analyte to mitigate sample losses and enhance specificity by providing prominent signal in both MS1 and MS2. Our approach includes downscaled and automated sample preparation in microliter volumes on 384-well plates for efficient, reproducible, and high throughput sample preparation, along with optimized reagent ratios for exhaustive labeling, and surface deactivated labware to further limit sample loss due to adsorption to surfaces. Critical parameters for Orbitrap sensitivity, such as RF voltage, HCD fragmentation energy, isolation window, MS2 accumulation time, and pAGC target, were specifically optimized for single-cell analysis using standards at typical single cell analyte concentrations (synthetic single-cell samples; see methods in the supplementary material). Moreover, the use of internal standards allows for the correction of possible differences in ionization suppression, retention time differences, and labeling efficiency. Integrating the reporter ion EICs on high-resolution MS2 provides additional specificity, in contrast to low mass resolution triple-quadrupole (QqQ) detection. Lastly, multiplexing enables high-throughput analysis of multiple individual samples simultaneously, significantly reducing analysis time.

However, the quantification is inherently constrained by the long scan durations required to achieve sufficient resolution for TMT10plex reporter ion isotopologues, extensive injection times to enhance sensitivity, and the objective to maintain short chromatographic peak widths to concentrate the analytes during elution. The analytes not retained here demonstrated also the lowest retention time in our previous work with similar column chemistry and UHPLC^42^, and the retention times of retained analytes showed a high correlation between this work and our previous work (retention time comparison in Figure S2, R>0.99). This suggests that the more polar analytes were not retained during the nLC operation steps prior to analytical gradient, such as 10 µL sample loading or gradient preparation. Also, an excessive reagent-to-analyte ratio was used to obtain efficient and exhaustive labeling, resulting in considerable hydrolyzed reagent peak (accurate *m/z* 248.1803) that limited the measurement to one tenth of the multiplexed single cell sample amount. Based on this, downscaling and reagent ratio optimization are among the most important parameters to advance isobaric labeling-based single cell metabolomics.

### Single cell analysis

The absolute analyte quantitation in single cells was performed by measuring the total of 60 individual HEK-293, TYK-nu, and TYK-nu.CP-r cells. The cells were sorted with CellenONE, with the simultaneous measurement of cell diameter, enabling the calculation of cell volume and thereby cellular analyte concentration. Six individual single cells were multiplexed with carrier, blank, and IS-blank. An example of MS2 reporter ion EIC of analytes from a single cell as well as internal standard is shown in Figure 5. The absolute analyte amounts were determined with the targeted method described above, and the multiplexed LC-MS2 results were parsed to individual cellular analyte amounts, as shown in Figure 6.

**Figure 5.**
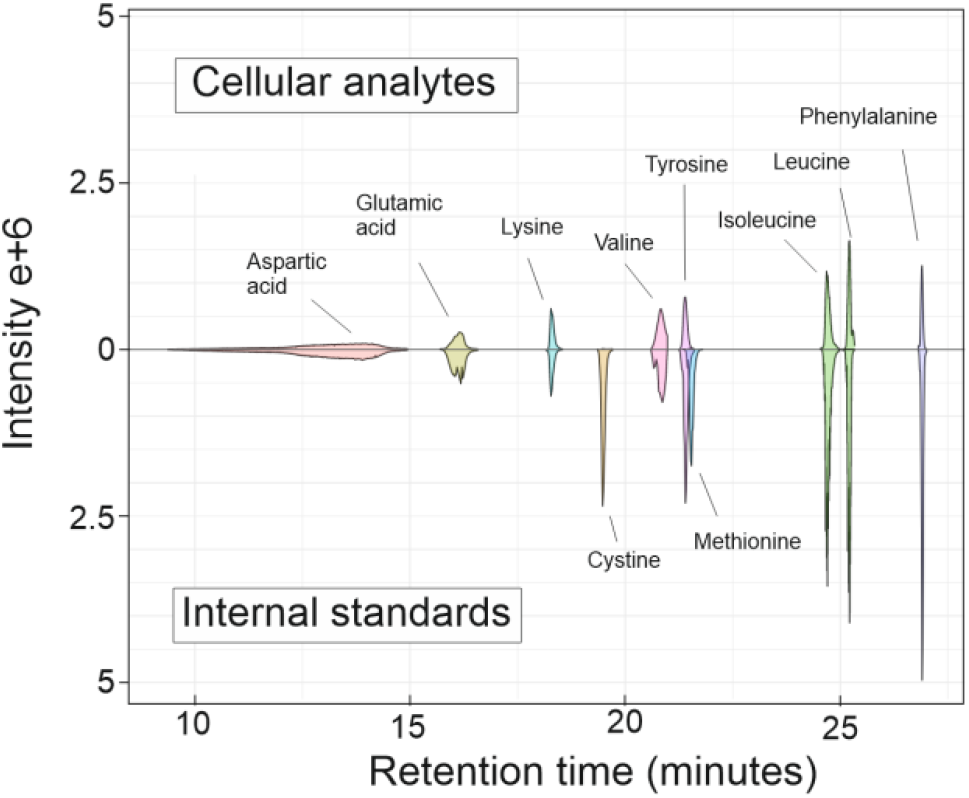
LC-MS2 integrated EICs of TMT10plex labeled amino acids in a single TYK-nu.CP-r cell (ID65). and used 12 fmol internal standard (EICs inversed).

**Figure 6.**
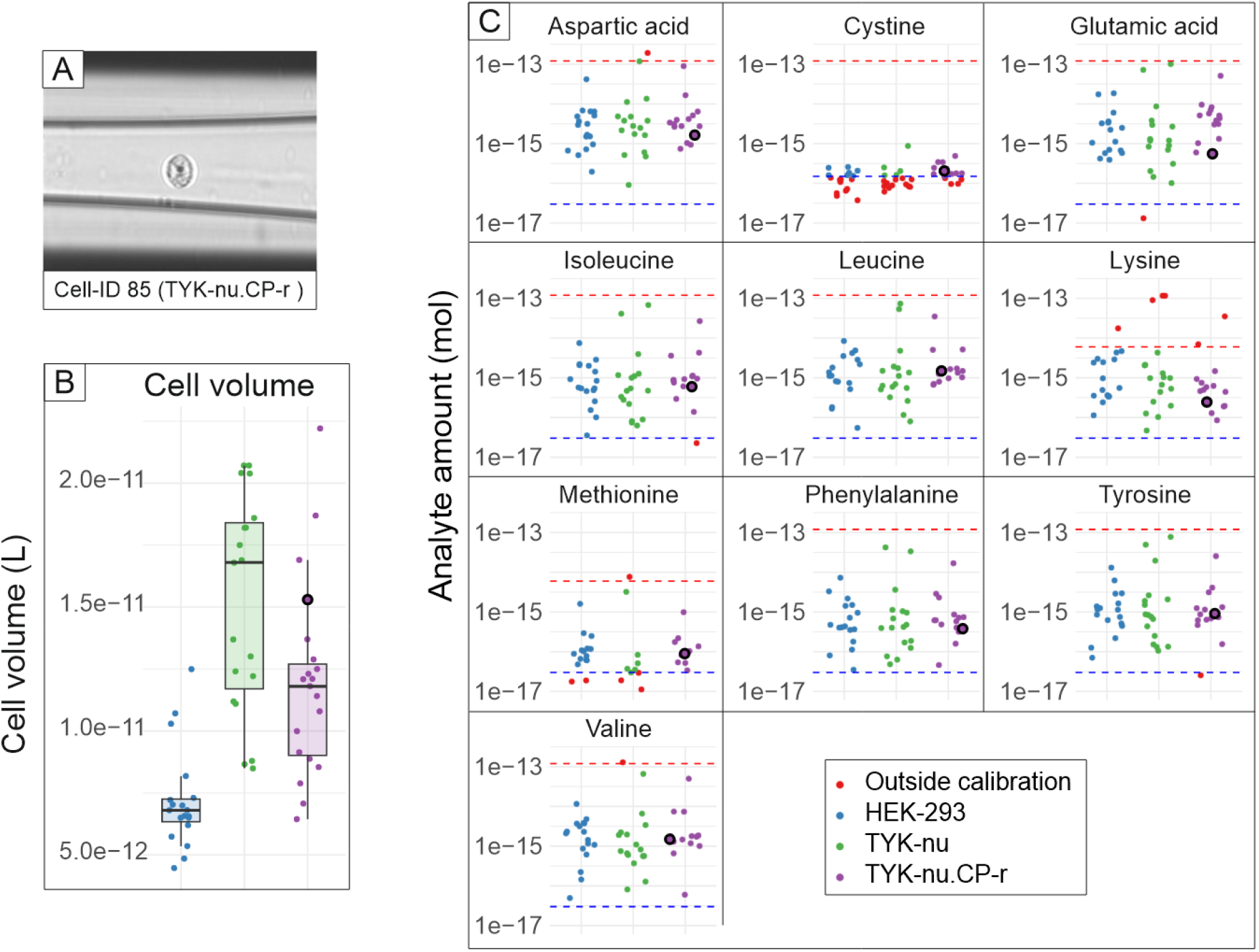
A) An example of CellenONE image shown with a TYK-nu.CP-r cell (ID85) attained during cell sorting. B) cell volume distribution of sorted cells by cell type with cell ID85 highlighted. C) Absolute amounts of ten amino acids in single cells along with the analyte lower limit of quantitation (LOQ, blue dashed line) and upper limit of quantitation (red dashed line) as in calibration. Calculated analyte amounts outside the scope of calibration are excluded (marked red) whereas the cell ID85 shown in panel A is highlighted.

The absolute cellular analyte amounts range from picomoles to femtomoles per cell, glutamic acid and aspartic acid being among the most abundant. Most measured analytes are well within the calibration range. However, the low amount of methionine and cystine, (that is mainly reduced to cysteine in the cytosol^49^) is notably low, resulting in cells with analyte amounts below LOQ (Figure 6).

This is also apparent on the single cell chromatogram in Figure 5, where single cell cystine and methionine are diminutive compared to the 12 fmol of heavy isotope internal standard. The cellular amounts are in the range that has been reported for HEK-293 cells.^55^ As the diameters of all individual cells and the absolute amounts of analytes in the same cells are known, cellular concentrations can be determined for analytes within calibration range. The intracellular concentrations are illustrated in Figure 7. On average, the TYK-nu cells were the largest, with cell volumes from 15.3 to 20.7 pL with an average of 15.3 pL, followed expectably by TYK-nu.CP-r with volumes from 6.4 to 22.2 pL and an average of 12.3 pL, whereas HEK-293 cells were approximately half of these, with volumes from 4.5 to 12.5 pL and an average of 6.7 pL. The cellular concentrations of amino acids in the measured cells are typically in the high micromolar range, with glutamic acid and aspartic acid occasionally reaching millimolar concentrations, similar to those reported for HEK-293 cells.^55^ Levene’s test indicated equal variances for most analyte concentrations between cell types.

**Figure 7.**
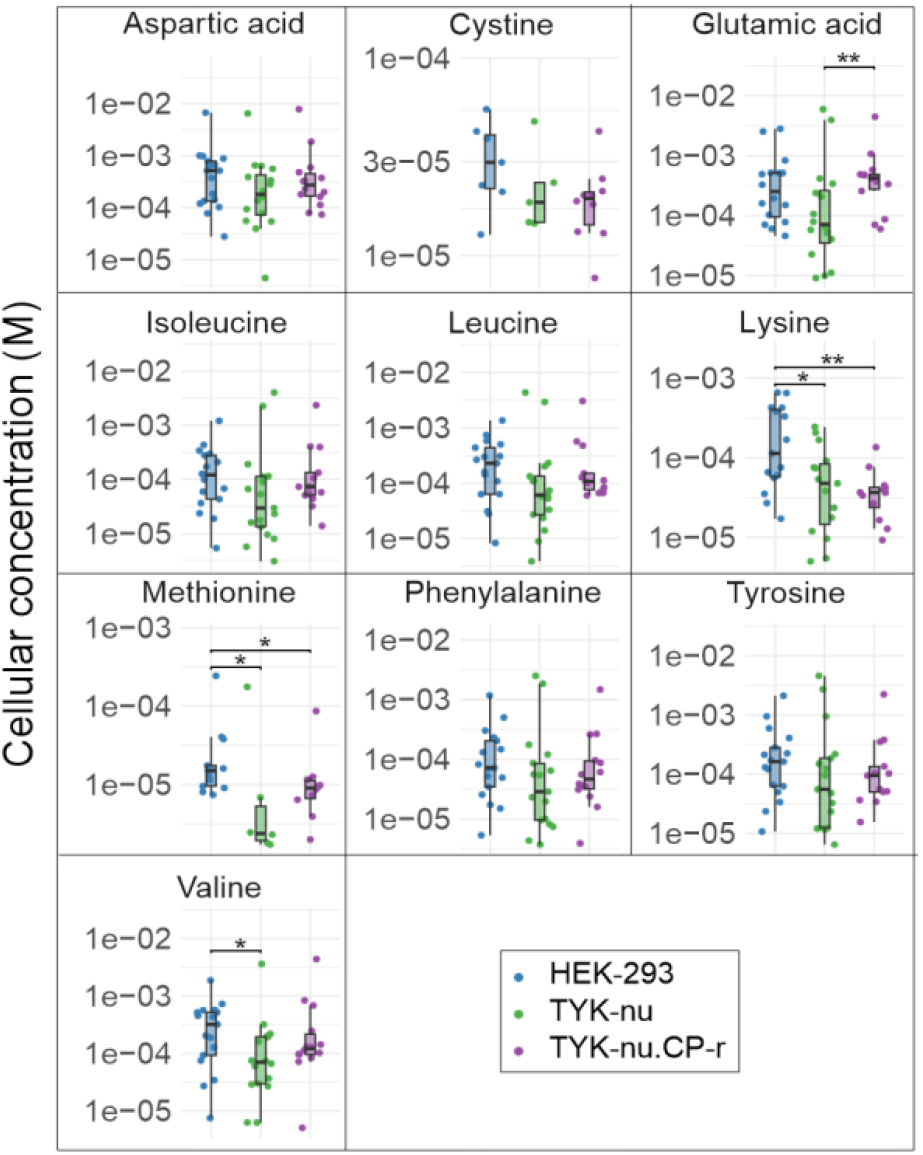
Cellular concentrations (mol/L) of amino acids in individual HEK-293, TYK-nu, and TYK-nu.CP-r cells based on absolute sample amount and cell size. The significance of 0.05 (*) and 0.01 (**) by one-way ANOVA.

Although the method demonstrated high repeatability at 3 fmol level, evidenced by a relative standard deviation (RSD) of 4.9%, there is a strikingly broad range in both absolute analyte quantities and cellular concentrations, extending across several orders of magnitude. The extent to which this reflects real biological effects remains uncertain. Possible errors may arise from factors such as inaccuracies in cell size determination, variation in sample preparation, inherent data processing sensitivity of multiplexed sample isotope purity effects and corrections, and the lack of cell cycle synchronization in cell culture. Nevertheless, the cell volumes, as illustrated in Figure 6B, do not exhibit large variations, with all being within a 3.5-fold ratio in cell volume. Cell sorting with CellenONE was performed on washed cells, with the acoustic dispensing itself resulting only in minuscule additional liquid volume. A similar freeze-thaw lysis method has been efficiently employed for single cells^56^, and the automated sample preparation is relatively robust highlighted by below 29% RSD in raw peak area of the 12 fmol IS analytes across the single cell samples. Additionally, the measured single cell analyte amounts are mostly well above the limit of quantification (LOQ). Therefore, it is unlikely that the observed significant differences are predominantly artificial.

Although considerable variation exists in the measured concentrations from cell to cell, statistically significant differences between cell groups were observed with glutamic acid, lysine, methionine, and valine. Generally, HEK-293 cells had higher analyte concentrations than TYK-nu or TYK-nu.CP-r cells, highlighting the similarity between the TYK-nu and TYK-nu.CP-r cell types. Only glutamic acid showed a statistically significant difference between TYK-nu and TYK-nu.CP-r cells. This may be attributed to the NADPH-glutathione (GSH) pathway, involving glutamate, as TYK-nu.CP-r cells’ resistance adaptation to the possibility of increased oxidative stress induced by cisplatin.^57^ It is possible that this pathway was active in the cells, although they were not challenged with cisplatin in this experiment.

The limited coverage of analytes and the relatively small number of single cells analyzed, combined with not challenging the cells with chemotherapy, restricts further elucidation of chemoresistance mechanisms in TYK-nu and TYK-nu.CP-r cells. A notable difference includes methionine being found at lower concentrations in TYK-nu variants compared to HEK-293 cells. Cancer cells, such as TYK-nu and TYK-nu.CP-r, may exhibit so-called methionine dependence, primarily participating in antioxidant defense through glutathione synthesis and serving as a precursor for S-adenosylmethionine, which is involved in the methylation of DNA, proteins, and lipids.^58^ As the metabolic flux and demand were not determined, the differences in cellular concentration could reflect either cancer adaptations or intrinsic differences between cell types.

### Untargeted cellular amine metabolomics

Untargeted single cell amine metabolomics was studied using 48 individual cells measured in three injection replicates. Untargeted feature detection and alignment were done on MS1 level using mzMine 3, whereas the quantitation was performed with TMT10plex reporter ions parsed from feature specific MS2 scans using a custom Python script. Orbitrap accumulation controlled by predictive automated gain control (pAGC) provides data that is inherently compositional^59^, requiring intra-scan normalization of MS2 scans that can be done with the carrier signal. However, the use of 127N blank and 127C IS-blank as feature selection controls is not feasible, as they suffer from carrier signal interference (also addressed as ion-coalescence) when large ratios are used, as reported previously.^60–63^

Untargeted metabolomics resulted in over 2000 features with accurate mass TMT10plex reporter ions in MS2 scans. These were filtered to 249 features (Figure S3). The loss of over 90% of features in filtering relates to the unresolved issue of high variance and missingness in single cell omics.^64,65^ In practice, there is the requirement for the same feature across samples to be selected during the moderately stochastic DDA selection for resolving the reporter ions connected to single cells. The cells with filtered features were projected on their principal components PC1 and PC2 (Figure 8).

**Figure 8.**
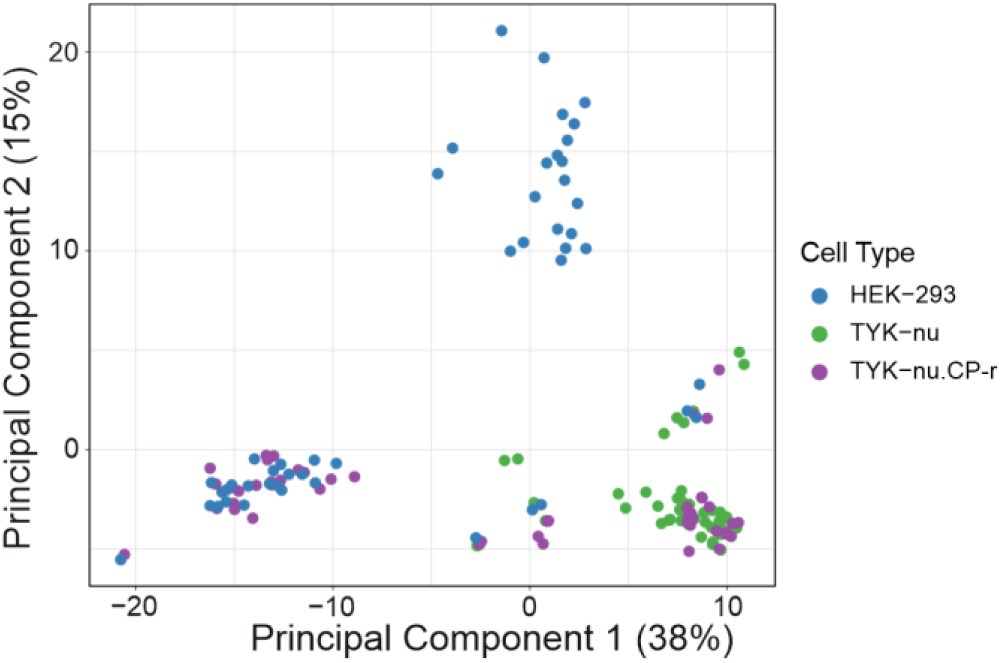
individual cells of untargeted amine metabolomics projected to their principal components and colored by cell type.

The PCA shows moderate cell line specific separation where the TYK-nu and TYK-nu.CP-r cluster closer to each other than HEK-293.

To elucidate the features contributing the most to this differentiation, we employed greedy forward feature selection with regularized least squares (RLS)^66^. Greedy RLS initiates with an empty feature set, and during each iteration it selects and adds the feature that yields the best performance based on leave-one-out cross-validation (LOOCV). To prevent selection bias, the effectiveness of the feature selection method was assessed using the standard nested cross-validation technique. Leave-cell-out was used in the outer-loop to count for the dependencies in sampling as there were three replications of each individual cell. Here, the feature selection was performed independently during each iteration of the outer cross-validation. The average performance of these models was then determined based on their one-vs-all prediction accuracy on the out-of-sample data across all rounds of the outer cross-validation, resulting in averaged accuracy of 0.59, indicating some classification power between the cell types.

A set of 100 features was selected using greedy RLS (File S1) where the selected features were first annotated using accurate MS1 and MS2 masses. Matching MS2 with TMT10plex labeled amines in our previous research^42^ provides high confidence annotation (Level 2), whereas using unlabeled MS2 fragment data in open metabolite libraries, such as GNPS, combined with TMT10plex labeling mass change is challenging due to different fragmentation of labeled analyte and fragmentation differences between CID and HCD, and could not be used to obtain more reliable identification (examples in Figure S4). Seven features could be annotated on Level 2 identification with relative intensities shown in Figure 9. From the resulting 93 features, further 44 features were putatively annotated using accurate precursor masses without the TMT10plex mass change against exact masses of HMDB library filtered to nitrogen containing compounds, resulting in Level 4 identification.

**Figure 9.**
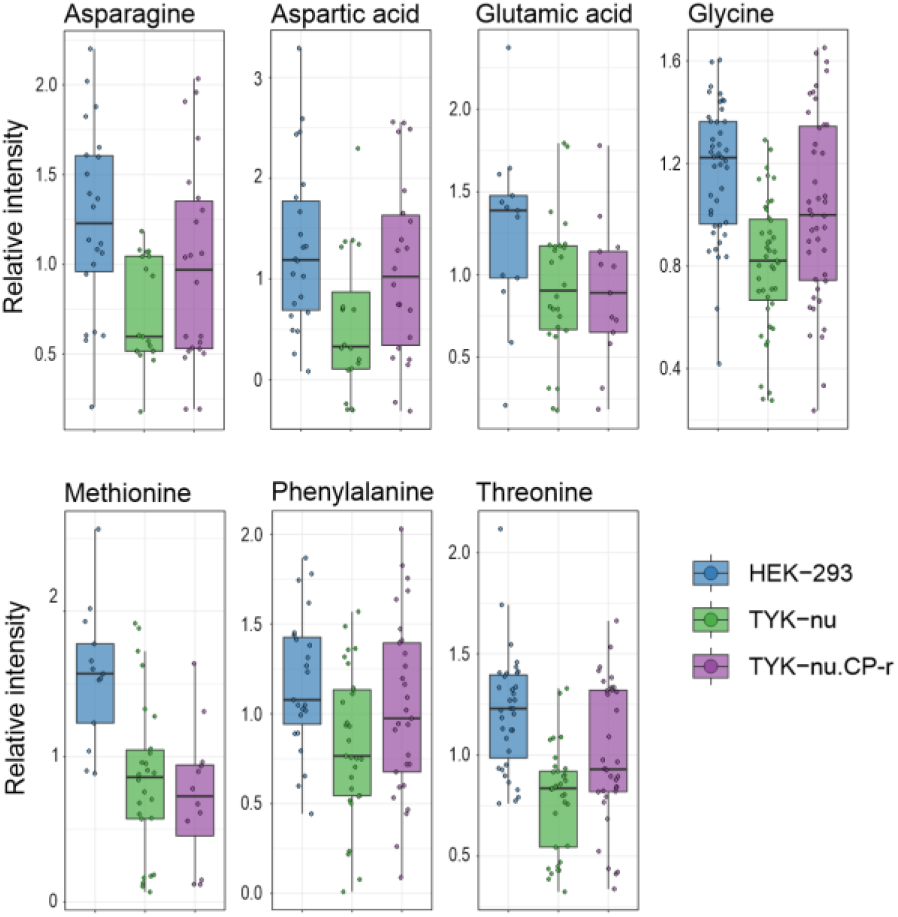
Metabolite comparison of confidently identified (Level2, matched to analyte standard MS2) analytes from Greedy RLS sub selection of untargeted metabolomics.

Amino acids play a crucial role in distinguishing between cell types, showing similar trends in both absolute and targeted quantification approaches, such as the lower methionine abundance in TYK-nu cell variants than HEK-293 (Figures 7 and 9). The untargeted approach highlights also the relatively broad range of analytes across individual cells.

Untargeted metabolomics using multiplexing and carrier offers a high-throughput and accessible approach for single-cell analysis. By increasing sample size and addressing ongoing challenges—such as manual data processing, the absence of comprehensive MS2 libraries of labeled analytes for annotation, and reproducible precursor selection during LC peak apex—we can gain even more profound insights into cellular metabolite differences.

## Conclusion

While proteomic approaches for single-cell analysis have emerged, metabolomics has not seen comparable advances, and single-cell metabolite analysis largely relies on relative quantitation obtained using specific or highly customized equipment and manual operation, resulting in limited method accessibility and low sample throughput. Here, we introduce an absolute nLC-HRMS2 quantitative approach for amines in single cells utilizing a multiplexing scheme with isobaric labeling, carrier concept, and automated, low-volume sample preparation. As demonstrated, this method can be applied to single cells, for both absolute quantitation of analytes in a targeted manner and relative quantitation of analytes in an untargeted manner. The method achieves high throughput of six multiplexed samples simultaneously, LOQs of low tens of attomoles, calibrated linear range typically from 30 amol to 120 fmol, and below 5% RSD for 3 fmol sample load, representative of the millimolar concentrations of amino acids typically found in single mammalian cells.

We applied the method to measure absolute amino acid concentrations and untargeted relative quantitation of amine analytes in individual cells of TYK-nu, TYK-nu.CP-r, and HEK-293 cell lines. Millimolar amino acid concentrations were determined, and we found concentration differences in lysine, methionine, and valine between HEK-293 against the TYK-nu variants, as well as difference in glutamic acid concentrations between the TYK-nu variants. In addition, we observed large cell to cell concentration differences as well as differences between cell types.

Although our approach is confined to amine analytes, amines take part in most metabolic processes, such as nucleotide metabolism, cofactor/redox metabolism, amino acid metabolism, energy metabolism, and acetyl group/carnitine metabolism. Metabolic pathways such as glycolysis/PPP, sugars, lipids, and TCA cycle metabolism, which primarily involve non-amine metabolites, could potentially be explored in single cells using a similar concept with existing isobaric reagents designed to target carboxylic acids^67^ or thiols^68,69^.

Instead of relying on relative quantities, quantifying the exact concentrations in individual cells can lead to a better understanding of the variability and distribution of cellular responses, processes, and disease states, such as cancer. Furthermore, absolute quantification provides comparability across different studies and could advance targeted therapeutic strategies and robust biomarker development. The utilization of the developed methodology can be extended to a wider range of other analytical applications that require high sensitivity.

## Supporting Information

The following data is provided as supporting information with additional references.^[70-91]^

- Supplementary data:
  ∘ Materials and methods
    ▪ Standards, cells, and reagents
    ▪ Single cell sample preparation
    ▪ Validation and optimization sample preparation
    ▪ LC-MS1 and LC-MS2 analysis
    ▪ Data processing
  ∘ Table S1. Labeled analytes and internal standards.
  ∘ Table S2. TMT exact masses and reporter ions
  ∘ Table S3. MS2 transition windows and precursors of the targeted tPRM method.
  ∘ Table S4. Exclusion mass list.
  ∘ Table S5. mzMine batch process.
  ∘ Figure S1. Calibration Graphs
  ∘ Figure S2. Retention time comparison on nLC and UHPLC.
  ∘ Figure S3. Heatmaps of single cells.
- Figure S4. Example of MS2 scan matching for glutamic acid annotationplementary file S1Sup: untargeted amine metabolite annotations.

## Acknowledgements

The authors would like to thank FIMM Single-Cell Analytics unit supported by HiLIFE and Biocenter Finland for CellenONE services, and Drug Discovery, Chemical Biology as well as the pipetting robot services with the facilities and expertise of the DDCB Unit at the Faculty of Pharmacy, University of Helsinki, supported by HiLIFE and Biocenter Finland. The authors would like to thank Dr. Nina Sipari for providing the use of sample preparation equipment. Research Council of Finland (project #321472) is acknowledged for funding.

