## Supplementary data for "Targeted and Untargeted Amine Metabolite Quantitation in Single Cells with Isobaric Multiplexing"

### Materials and methods

#### Standards, cells, and reagents

*Reagents.* Amino acid standards and the corresponding amino acids with heavy $^{15}N$ and $^{13}C$ isotopes (Cambridge Isotope Laboratories) were used as internal standards (Table S1). These are referred to as light and heavy amino acid standard, respectively. LC-MS grade water and acetonitrile, both with 0.1% formic acid, were from Thermo Scientific. TMT0 and TMT10plex isobaric reagents, 1 M triethyl ammonium bicarbonate (TEAB) buffer, and 50% hydroxylamine solution were purchased from Thermo Scientific. Anhydrous dimethyl sulfoxide (DMSO) was acquired from Honeywell.

*Cell culture, imaging, sorting, and lysis*. TYK-nu and TYK-nu.CP-r cell lines were a kind gift from Jukka Saarinen, and the HEK293 cell line was received from Leena Pietilä (both from University of Helsinki). All cell lines tested negative for mycoplasma at the beginning of cell culture. The cells were grown at 37 °C under 5% $CO_2$ in antibiotic-free low glucose Dulbecco's Modified Eagle Medium with L-glutamine, pyruvate, and HEPES (Gibco, REF 22320022), supplemented with 10% fetal bovine serum (Gibco, REF 10270106). At >70% confluence, the cells were washed with phosphate-buffered saline (PBS; Gibco, REF 14200-067, diluted into 1X), detached using TrypLE Express (Gibco, REF 12604021), washed twice with cold PBS, and resuspended in cold PBS. The cells were kept on ice until sorting on the same day. Single cells were individually imaged and sorted using the cellenONE platform into 384-well plate wells containing 5 µL of water. Circularity, diameter, and elongation cut-off values were used to exclude doublets and debris. Sorted and imaged cells were frozen in -80 °C until cell lysis. During sorting, the cell viability was >85%, as assessed with LUNA-FL automated cell counter. For the samples prepared in bulk, cells were sorted with flow cytometry using BD FACSAria II. Cell lysis was done one plate at a time by a freeze-thaw lysis approach modified from that presented by Végvári *et al.* [1]. Cell lysis consisted of four 2-min cycles of freezing in liquid nitrogen, with thawing in 37°C between the freezing cycles. Next, the cells were sonicated for 5 min and heated to 90 °C for 10 min. Finally, the lysates were centrifuged to the bottoms of the wells, and the 384-well plates were sealed using well plate sealing foil.

#### Single cell sample preparation

Multiplexed samples consisted of approximately 400 cell-equivalent (120 fmol) carrier (labeled with TMT10plex label 126) of either analyte standard for targeted approach or cell lysate for untargeted approach, blank (127N), blank with internal standards of heavy amino acids (IS-blank, 128N), and six single cells (128C, 129N, 129C, 130N, 130C, and 131; exact masses of reporter ions in Table S2). TMT10plex label 127C was not used due to isotopic interference from the carrier, caused by isotopic impurity in the labeling reagents. The labeling principle is the same as previously reported.[2]

**Carrier for targeted metabolomics** was prepared by mixing 79 µL of 100 mM TEAB, 180 µL of a solution containing 240 nM of each light amino acid standard in water, and 4.5 µL of 960 nM heavy amino acid standard in water, in a low-binding Eppendorf tube. This was labelled using 12 µL of 17.2 mM TMT10plex 126 label reagent dissolved in anhydrous DMSO, and allowed to react for one hour. Subsequently, the labelling reaction was quenched with 12 µL of 1% hydroxylamine, and this mixture was incubated for 15 minutes.

**Carrier for untargeted metabolomics** was prepared similarly as for targeted metabolomics, but using 180 µL of 80 cells/µL lysate in water instead of the light amino acid standard.

**Blank** was prepared using 260 µL of TEAB, labeled with 127N label reagent as described above. **IS blank** was produced by mixing 353 µL of 100 mM TEAB with 6 µL of 960 nM heavy amino acid standard, and labeled with 128N label reagent as described above.

The lysed **individual real single cell samples** on the 384-well plates were prepared using a Biomek i7 pipetting robot. 2 µL of 60 nM heavy amino acid standard (internal standard) was added into each well containing a lysed single cell, followed by labeling with 1 µL of one of the TMT10plex reagents (128C, 129N, 129C, 130N, 130C, or 131). To limit batch effects, label swap was carried out, so that the same cell line was not always labeled using same reagent. Labeling was continued for 1h in RT, and the labeling was then quenched using 1 µL of 1% hydroxylamine.

60 lysed and labeled individual single cells were used for targeted and 48 for untargeted metabolomics. Cell samples were combined in sets of six individual cells (labeled with one of each reagent from 128C to 131), along with 8 µL of the corresponding carrier, blank, and IS-blank into low binding 250 µL glass vial inserts. The samples were evaporated to dryness with a vacuum centrifuge and frozen to -80 °C until measurement. Prior to analysis, the samples were thawed and dissolved in 20 µL of eluent A used in nLC.

#### Validation and optimization sample preparation

MS2 sensitivity specific parameters were optimized using a synthetic single cell TMT10plex labeled sample in larger volume providing comparable measurements. This synthetic single cell sample consisted, similarly to real single cell samples, of carrier, blank, IS-blank and six single cell equivalent samples. Absolute quantitation was obtained using validation samples for calibration, linearity, LOD, LOQ, and precision.

**Synthetic single cell sample**, used for optimizing MS2 sensitivity specific parameters, was prepared so that one LC-MS injection of it corresponds to the analyte amount in one multiplexed single cell sample. 40 µl of 100 mM TEAB, 10 µl of 6 nM light amino acid standard, and 40 µL of 60 nM heavy amino acid standard were added into six separate low-binding Eppendorf tubes. Each of these was labeled with one TMT10plex isotopomer (128C-131, 17.2 mM in DMSO) for 1h and quenched with 20 µL of 1% hydroxylamine for 15 min. After labeling, the six synthetic single cell samples were combined with 100 µL of carrier for targeted metabolomics, 100 µL of blank, and 100 µL of IS-blank sample, and evaporated to dryness. The sample was then dissolved in 200 µL of eluent A.

**TMT0-labeled sample**, used for optimizing the general LC and MS parameters, was prepared in a low-binding Eppendorf tube by combining 120 µL of 100 mM TEAB, 750µL of 240 nM light amino acid standard, and 300 µL of 204 nM heavy amino acid standard. This was labeled with 60 µL of 17.2 mM of TMT0 in DMSO for 1h, and quenched with 60 µL of 1% hydroxylamine. The sample was evaporated to dryness and dissolved into 600 µL of eluent A, and further diluted 1:1, 1:10 and 1:100 with eluent A.

**Validation samples** followed the same multiplexing framework, and they were prepared individually from stock solutions into 384-well plates using the Biomek i7 pipetting robot. For each sample, 10 µL of TEAB, 4 µL of 60 nM heavy amino acid standard, and varying amounts of light amino acid standard were added to obtain final sample amounts from 0.03 fmol to 120 fmol of light amino acids per sample injection. These samples were used for calibration, linearity, LOD, and LOQ, as well as multiple 0.3 fmol and 3.0 fmol sample amounts for precision (method repeatability). Samples were labeled with the appropriate TMT10plex isotopomer for 1h and quenched with 2 µL of 1% hydroxylamine for 15 minutes. The individual samples were then multiplexed and dried using a vacuum centrifuge. The validation samples were dissolved in 40 µL of eluent A before measurement.

#### LC-MS1 and LC-MS2 analysis

The LC-MS measurements were carried out using an Orbitrap Fusion tribrid mass spectrometer coupled to an EASY nLC 1200 liquid chromatograph with a nanospray Flex ion source (all from Thermo Fisher Scientific).

Chromatography was performed using Waters Acquity M class C-18 column (M-Class HSS T3 1.8 µm 75 µm x 200 mm) at room temperature, and the column was connected to a stainless steel nano bore emitter (30 µm I.D. 40 mm, Thermo Scientific) using a MicroTight union (PEEK 1/32" P-771, IDEX Health & Science). Eluent A consisted of 0.1% formic acid in water, and eluent B consisted of 0.1% formic and 80% acetonitrile in water. The flow rate was 300 nL $min^{-1}$. The gradient started from 1% eluent B and was then increased to 20% B in 20 min, to 95% B in the next 10 min, and held at 95% B for 20 min. The gradient was then decreased to 1% B in 1 min and held at 1% B for 9 minutes. The resulting total gradient time was 60 min. After each injection, the autosampler was washed with 28 µL of eluent A, 28 µL of eluent B, and again with 28 µL of eluent A.

Targeted tPRM windows were built with MS1 measurements of optimization samples using orbitrap with 120k resolution, *m/z* 100-1600 scan range, and 100 ms maximum injection time with 250% normalized AGC target. RF lens was optimized to 80%, spray voltage to 2.1 kV, sweep gas flow to 0.5 arbitrary units, and ion transfer tube temperature to 250 °C. The sweep cone was not installed, and the low-flow pure nitrogen sweep gas (not directed at the spray) was mainly used to keep contaminants in the room air from entering the open ion source and to stabilize the electrospray. The targeted MS2 measurements were performed using timed parallel reaction monitoring (tPRM) and timed precursor isolation based on analyte retention times measured with the MS1 method (tPRM windows in Table S3). Internal mass calibration with Easy-IC (fluoranthene) was used in the beginning of each measurement (RunStart EasyIC). MS2 precursor isolation was done using quadrupole mass window of 1.0 Da, and the higher-energy collisional dissociation (HCD) normalized collision energy was optimized to 35 % and used for all analytes. Reporter product ions were detected with a mass resolution of 60 000 and automatic scan range with defined low mass *m/z* 110 to ensure the detection of TMT reporter ions. AGC target was optimized to 250% with a maximum injection time of 150 ms.

For untargeted amine metabolomics, the analysis was done using data dependent acquisition (DDA) approach. The chromatographic gradient was adjusted to achieve 40% B instead of 20% at 20 min, but all other LC parameters were kept similar as with the tPRM method. Spray voltage was set to 2.0 kV, sweep gas at 0.5 arbitrary unit flow and ion transfer tube to 250 °C. Internal mass calibration was done using RunStart Easy IC. MS scan (survey scan) consisted of orbitrap analysis with scan range *m/z* 100-1600, 120 k resolution and with wide quadrupole isolation enabled. The maximum injection time was set to 50 ms with normalized AGC target of 250%. RF lens was set to 80%. The data dependent MS2 scans were based on top 7 precursors with targeted exclusion mass list (Table S4) of background ions determined based on a blank injection, intensity threshold of 3E4, and 20 s dynamic exclusion with 10 ppm mass tolerance. Also, apex detection filter of 4 s FWHM (based on MS1 optimization samples) and desired window of 40% was used. MS2 scans were done using orbitrap with 60 k resolution and 1 Da isolation window. Scan range was set to begin from 110 Da to include the TMT reporter ions, and HCD fragmentation with 37% normalized collision energy was used. Maximum ion injection time was set to 118 ms with normalized AGC target to 250%.

#### Data processing

Raw and metadata are provided in MassIVE repository with identifier MSV000095047 and password SCM2024. For targeted metabolite analysis, raw tPRM MS2 data was imported to Skyline[3] (22.2.1.425), along with the analyte to reporter ion transition list. Peaks in the MS2 reporter ion chromatograms were integrated, and the results were exported to Rstudio for further analysis (Rstudio[4] version 2023.09.1 Build 494 with R[5] version 4.2.2). In R, data processing and visualization were done using the Cran package "tidyverse"[6] including “dplyr” and “tidyr”. Raw mzML data was processed with “MSnbase”[7], and the graph zoom visualization was implemented using “ggforce”[8] and “scales”[9] and the color scheme was controlled using “RColorBrewer”[10]. The code for the full data analysis is in Table S3. Briefly, isobaric interference between reporter ion signals was corrected using inverse matrix multiplication and the isotopic purity of the label reagents provided in the Certificate of Analysis of the TMT reagent batch.

The targeted method was partially validated according to ICH guidelines[11] in terms of specificity, linearity, limit of quantitation (LOQ), and method repeatability. LOQ was determined as the lowest measured concentration producing a signal to noise ratio (S/N) > 10. Because MS2 noise within 5 ppm of TMT fragment ions is mainly not present, LOQ was further defined to have a relative residual between calibration line and actual measurement value below 60%. Analyte signals corrected for isotopic interference were calculated as relative to the corresponding corrected internal standard signals, and the absolute analyte amount in sample was calculated using calibration line with calibration sample concentrations. The calibration lines were produced using weighed (1/concentration) linear regression, and they were evaluated by $R^2$ as well as residuals of measurements against calibration line. Within range outliers were determined based on the residuals using Grubbs outlier test based on both residuals and relative residuals. Calibration sample 0.3 fmol was outlier with all analytes and 0.6 fmol with some analytes. Seven technical replicates at 0.3 fmol and 3 fmol per injection were measured for method repeatability. One technical replicate of 0.3 fmol was also tested to be an outlier and removed from all analytes.

The absolute sample amounts in single cells were calculated similarly using the calibration line equations. Cell volumes were calculated as spheres using diameter as elongation and circularity were used for cut-off values the measured absolute sample amounts were divided by the cell volume to obtain cellular analyte concentration. The homoscedasticity of cellular concentrations was tested with Levene's test for equal variance and Kruskal-Wallis one way analysis of variance was used to compare the different analytes between cell types followed by Tukey's HSD post-hoc test.

For the untargeted amine metabolomics, Thermo raw data was converted to mzML using MSconvert[12] (3.0.22238) and imported to MZmine[13] 3 (version 3.9.0). The files were batch processed to feature table of features including MS2 scan (MzMine batch process is in Table S5) Data processing and analysis were performed using Python 3.11.3 in Visual Studio Code (version 1.89.1). The data was processed as data frame using the Pandas library, with numerical operations facilitated by NumPy[14]. Statistical computations, including standardization and normalization, cosine similarity and dimensionality reduction utilized functions from the SciPy[15] and scikit-learn[16] libraries. Batch effects were corrected using PyComBat[17], a Python implementation of the ComBat[18] algorithm. Visualization of the processed data was achieved through the use of Matplotlib[19] and Seaborn[20].

The absolute TMT10plex reporter ion intensities were parsed from MS2 scans and normalized to the carrier reporter ion intensity. Features were filtered to those detected in at least 25% of the cells, and accurate mass over 250 Da. To take into account the nLC retention time differences between samples, features within 0.005 Da and 0.4 min were binned, and features within bins were filtered to most abundant. Labeling byproducts were removed and the missingness was filled the smallest intensity, and the data was log2-transformed and Z-score standardized. Batch effects between multiplexed samples were corrected using ComBat, and the measurements were projected to their principal components PC1 and PC2.

The features driving the separation on PCA were inspected using supervised classification Greedy forward feature selection with regularized least squares (RLS) from Rlscore Library[21]. Greedy RLS initiates with an empty feature set, and during each iteration it selects and adds the feature that yields the best performance based on Leave-One-Out Cross-Validation (LOOCV). To prevent selection bias, the effectiveness of the feature selection method was assessed using the standard nested cross-validation technique where the feature selection was performed independently during each iteration of the outer cross-validation. The average performance of these models was then determined based on their one-vs-all prediction accuracy on the out-of-sample data across all rounds of the outer cross-validation. As there are three replications of each sampled cell, to count for the dependencies in sampling, leave-cell-out is used in the outer-loop.

The resulting features were searched against MS2 fragment spectra of TMT10plex labeled amines from our previous work[2] as well as MS1 against HMDB[22] library (downloaded in March 2024) filtered to amine containing chemical formulas with 5 mDa window. Feature MS2 annotation was considered also with MS2 spectra in open libraries in GNPS[23] with added TMT10plex modification (+229.162932 Da). Features were annotated using 5 level confidence hierarchy[24] to Level 2 using in-house high mass resolution MS2 data of TMT10plex labeled analytes and to Level 4 using the accurate mass precursors against nitrogen containing compounds of HMDB.

#### Table S1. Labeled analytes and internal standards.

Analytes, their measured ion ions exact masses of ions as well as internal standard and internal standard exact mass.

| **Amino acid** | **Ion** | **Exact mass** | **Internal standard** | **Exact mass** |
|---|---|---|---|---|
| Alanine | $[1M+1TMT10+1H]^+$ | 319.217893 | Alanine ($^{13}C_3$ $^{15}N_1$) | 323.224983 |
| Arginine | $[1M+1TMT10+1H]^+$ | 404.281893 | Arginine ($^{13}C_6$ $^{15}N_4$) | 414.290153 |
| Aspartic acid | $[1M+1TMT10+1H]^+$ | 363.207723 | Aspartic acid ($^{13}C_4$ $^{15}N_1$) | 368.218163 |
| Glutamic acid | $[1M+1TMT10+1H]^+$ | 377.223373 | Glutamic acid ($^{13}C_5$ $^{15}N_1$) | 383.237163 |
| Glycine | $[1M+1TMT10+1H]^+$ | 305.202243 | Glycine ($^{13}C_2$ $^{15}N_1$) | 308.205983 |
| Histidine | $[1M+1TMT10+1H]^+$ | 385.239693 | Histidine ($^{13}C_6$ $^{15}N_3$) | 394.250913 |
| Isoleucine | $[1M+1TMT10+1H]^+$ | 361.264843 | Isoleucine ($^{13}C_6$ $^{15}N_1$) | 368.281983 |
| Leucine | $[1M+1TMT10+1H]^+$ | 361.264843 | Leucine ($^{13}C_6$ $^{15}N_1$) | 368.281983 |
| Lysine** | $[1M+2TMT10+2H]^{2+}$ | 303.222973 | Lysine ($^{13}C_6$ $^{15}N_2$) | 307.230063 |
| Methionine | $[1M+1TMT10+1H]^+$ | 379.221263 | Methionine ($^{13}C_5$ $^{15}N_1$) | 385.235053 |
| Phenylalanine | $[1M+1TMT10+1H]^+$ | 395.249193 | Phenylalanine ($^{13}C_9$ $^{15}N_1$) | 405.276383 |
| Proline | $[1M+1TMT10+1H]^+$ | 345.233543 | Proline ($^{13}C_5$ $^{15}N_1$) | 351.247333 |
| Serine | $[1M+1TMT10+1H]^+$ | 335.212803 | Serine ($^{13}C_3$ $^{15}N_1$) | 339.219893 |
| Threonine | $[1M+1TMT10+1H]^+$ | 349.228453 | Threonine ($^{13}C_4$ $^{15}N_1$) | 354.238893 |
| Tyrosine | $[1M+1TMT10+1H]^+$ | 411.244103 | Tyrosine ($^{13}C_9$ $^{15}N_1$) | 421.271293 |
| Valine | $[1M+1TMT10+1H]^+$ | 347.249193 | Valine ($^{13}C_5$ $^{15}N_1$) | 353.262983 |
| Cystine** | $[1M+2TMT10+2H]^{2+}$ | 350.182133 | Cystine ($^{13}C_6$ $^{15}N_2$) | 354.189223 |

** Lysine and cystine are double labeled and doubly charged

### Table S2. TMT exact masses and reporter ions

Labeling exact mass change with TMT10plex reagent as well as reporter (fragment) ion exact mass produced by HCD.

| TMT reagent | Labeling mass change | HCD reporter ion exact mass |
|---|---|---|
| TMT10plex-126 | 229.162932 | 126.127726 |
| TMT10plex-127N | 229.162932 | 127.124761 |
| TMT10plex-127C | 229.162932 | 127.131081 |
| TMT10plex-128N | 229.162932 | 128.128116 |
| TMT10plex-128C | 229.162932 | 128.134436 |
| TMT10plex-129N | 229.162932 | 129.131471 |
| TMT10plex-129C | 229.162932 | 129.137790 |
| TMT10plex-130N | 229.162932 | 130.134825 |
| TMT10plex-130C | 229.162932 | 130.141145 |
| TMT10plex-131 | 229.162932 | 131.138180 |

Table S3. MS2 transition windows and precursors of the targeted tPRM method.

| **Compound** | **Precursor (m/z)** | **Precursor isolation window (min)** |
|---|---|---|
| Aspartic acid | 363.2077 | 9 - 16 |
| IS Aspartic acid | 368.2167 | 9 - 16 |
| Threonine | 349.2285 | 12.8 - 17.5 |
| IS Threonine | 354.2375 | 12.8 - 17.5 |
| Alanine | 319.2179 | 13.5 - 17.5 |
| IS Alanine | 323.2239 | 13.5 - 17.5 |
| Glutamic acid | 377.2234 | 13.9 - 18 |
| IS Glutamic acid | 383.2354 | 13.9 - 18 |
| Serine | 335.2128 | 14.5 - 18.5 |
| IS Serine | 339.2000 | 14.5 - 18.5 |
| Proline | 345.2335 | 16.9 - 21 |
| IS Proline | 351.2455 | 16.9 - 21 |
| Lysine | 303.2230 | 17 - 20 |
| IS Lysine | 308.2334 | 17 - 20 |
| Cystine | 350.1821 | 18.2 - 21 |
| IS Cystine | 354.1881 | 18.2 - 21 |
| Valine | 347.2492 | 19.8 - 23 |
| IS Valine | 353.2612 | 19.8 - 23 |
| Methionine | 379.2213 | 20 - 23.5 |
| IS Methionine | 385.2333 | 20 - 23.5 |
| Tyrosine | 411.2441 | 20.2 - 23 |
| IS Tyrosine | 421.2681 | 20.2 - 23 |
| Isoleucine | 361.2648 | 23.7 - 25 |
| IS Isoleucine | 368.2798 | 23.7 - 25 |
| Leucine | 361.2648 | 24.9 - 27 |
| IS leucine | 368.2798 | 24.9 - 27 |
| Phenylalanine | 395.2492 | 26 - 28.5 |
| IS Phenylalanine | 405.2732 | 26 - 28.5 |

### Table S4. Exclusion mass list.

An exclusion mass list was produced based on background ions in both A-eluent, B-eluent and labeling byproduct *m/z* values with relative intensity above 1% in mass spectra.

| **Number** | **m/z** | **Number** | **m/z** | **Number** | **m/z** | **Number** | **m/z** |
|---|---|---|---|---|---|---|---|
| 1 | 202.0773 | 21 | 280.1618 | 41 | 145.1220 | 61 | 229.1403 |
| 2 | 149.0230 | 22 | 217.1066 | 42 | 520.1378 | 62 | 315.2132 |
| 3 | 279.1585 | 23 | 221.1531 | 43 | 229.0009 | 63 | 301.1401 |
| 4 | 445.1187 | 24 | 223.0631 | 44 | 163.1114 | 64 | 339.3440 |
| 5 | 219.1738 | 25 | 209.0595 | 45 | 256.2629 | 65 | 215.1247 |
| 6 | 429.0875 | 26 | 136.0213 | 46 | 268.9778 | 66 | 360.3226 |
| 7 | 355.0690 | 27 | 447.1159 | 47 | 220.1772 | 67 | 354.3176 |
| 8 | 203.0806 | 28 | 209.1531 | 48 | 277.1792 | 68 | 273.1664 |
| 9 | 371.1002 | 29 | 150.0263 | 49 | 667.1746 | 69 | 441.2961 |
| 10 | 271.2625 | 30 | 371.3146 | 50 | 238.0129 | 70 | 149.0229 |
| 11 | 203.0850 | 31 | 235.2051 | 51 | 372.0999 | | |
| 12 | 235.1687 | 32 | 199.1689 | 52 | 163.0750 | | |
| 13 | 519.1373 | 33 | 170.0961 | 53 | 593.1560 | | |
| 14 | 220.1817 | 34 | 253.1792 | 54 | 357.0660 | | |
| 15 | 446.1189 | 35 | 231.0337 | 55 | 151.1114 | | |
| 16 | 503.1062 | 36 | 161.0957 | 56 | 202.0771 | | |
| 17 | 391.2832 | 37 | 229.0181 | 57 | 203.0804 | | |
| 18 | 205.0854 | 38 | 431.0845 | 58 | 338.3408 | | |
| 19 | 299.0610 | 39 | 359.0275 | 59 | 203.0849 | | |
| 20 | 430.0875 | 40 | 356.0686 | 60 | 271.1872 | | |

### Table S5. mzMine batch process.

Batch process steps in the mzMine for the untargeted metabolomics experiment results.

| Step | Description | Settings |
|---|---|---|
| **Data Import** | Imports raw spectral data from files. | **File Names**: List of .mzML files |
| **Mass Detection (MS1)** | Detects masses in the MS1 spectral level. | **Mass Detector**: Factor of lowest signal; **Noise Factor**: 2; **Denormalize Fragment Scans**: Enabled |
| **Mass Detection (MS2)** | Detects masses in the MS2 spectral level. | **Mass Detector**: Factor of lowest signal; **Noise Factor**: 1; **Denormalize Fragment Scans**: Enabled |
| **Chromatogram Building (ADAP)** | Constructs chromatograms from the detected masses. | **Minimum Consecutive Scans**: 4; **Minimum Intensity**: 2000.0; **Minimum Absolute Height**: 10000; **m/z Tolerance**: 0.01 absolute, 10.0 ppm |
| **Smoothing** | Applies smoothing algorithms to chromatograms. | **Algorithm**: Savitzky-Golay; **Polynomial Order**: 3 |
| **Local Minimum Resolver Deconvolution** | Resolves overlapping signals within chromatograms. | **Algorithm**: Local Minimum Resolver; **MS/MS Scan Pairing**: Enabled; **Dimension**: Retention Time; **Chromatographic Threshold**: 85%; **Minimum Search Range RT**: 0.3; **Minimum Relative Height**: 0; **Minimum Absolute Height**: 10000; **Min Ratio of Peak Top/Edge**: 1.8; **Peak Duration Range**: 0-9.01; **Minimum Scans**: 4 |
| **13C Isotope Grouper** | Groups isotopic peaks together. | **m/z Tolerance**: 0.01 absolute, 3.0 ppm; **RT Tolerance**: 0.1 minute; **Monotonic Shape**: Enabled; **Maximum Charge**: 2; **Representative Isotope**: Most intense; **Never Remove Feature with MS2**: Enabled |
| **Isotopic Peak Finder** | Identifies specific isotopic peaks within the MS data. | **Chemical Elements**: H, C, N, O, S; **m/z Tolerance**: 0.01 absolute, 3.0 ppm; **Maximum Charge of Isotope m/z**: 1; **Search in Scans**: Single most intense |
| **Feature Alignment (RANSAC)** | Aligns features across different samples. | **m/z Tolerance**: 0.005 absolute, 5.0 ppm; **RT Tolerance**: 3.0 minutes; **RT Tolerance After Correction**: 0.2 minutes; **RANSAC Iterations**: 0 (not limited); **Minimum Number of Points**: 50%; **Threshold Value**: 0.1 |
| **Feature Lists Rows Filter** | Filters the feature rows based on selections. | **Minimum Aligned Features (Samples)**: 1 or 0%; **Feature with MS2 Scans**: Enabled; **Never Remove Feature with MS2**: Enabled |
| **Peak Finder (Multithreaded)** | Identifies and fills gaps in the feature list. | **Intensity Tolerance**: 20%; **m/z Tolerance**: 0.01 absolute or 10 ppm; **Retention Time Tolerance**: 1.5 minutes; **Minimum Scans**: 1 |
| **Duplicate Filtering** | Removes duplicate features. | **Filter Mode**: New average; **m/z Tolerance**: 0.01 absolute, 2.0 ppm; **RT Tolerance**: 0.3 minute |
| **Correlation Grouping (MetaCorrelate)** | Groups features based on correlation analysis. | **RT Tolerance**: 0.33 minutes; **Minimum Feature Height**: 0; **Intensity Threshold for Correlation**: 2; **Feature Shape Correlation**: Enabled; **Feature Height Correlation**: Enabled |
| **Ion Identity Networking** | Constructs networks of ion identities based on mass spectral data. | **m/z Tolerance**: 0.01 absolute or 3 ppm; **Min Height**: 0; **Ion Identity Library**: As suggested; **Annotation Refinement**: Enabled |
| **MSMS Spectral Networking** | Links MS/MS spectra across samples to build spectral networks. | **m/z Tolerance (MS2)**: 0.01; **Only Best MS2 Scan**: Enabled; **Max Precursor m/z Delta**: 500; **Minimum Matched Signals**: 4; **Minimum Cosine Similarity**: 0.7 |
| **Data Export as CSV** | Exports the processed data to a CSV file for further analysis or reporting. | **Export All**: Enabled |

### Figure S1. Calibration Graphs

Calibration graphs of partially validated (as in article table 1) analyte quantitation.

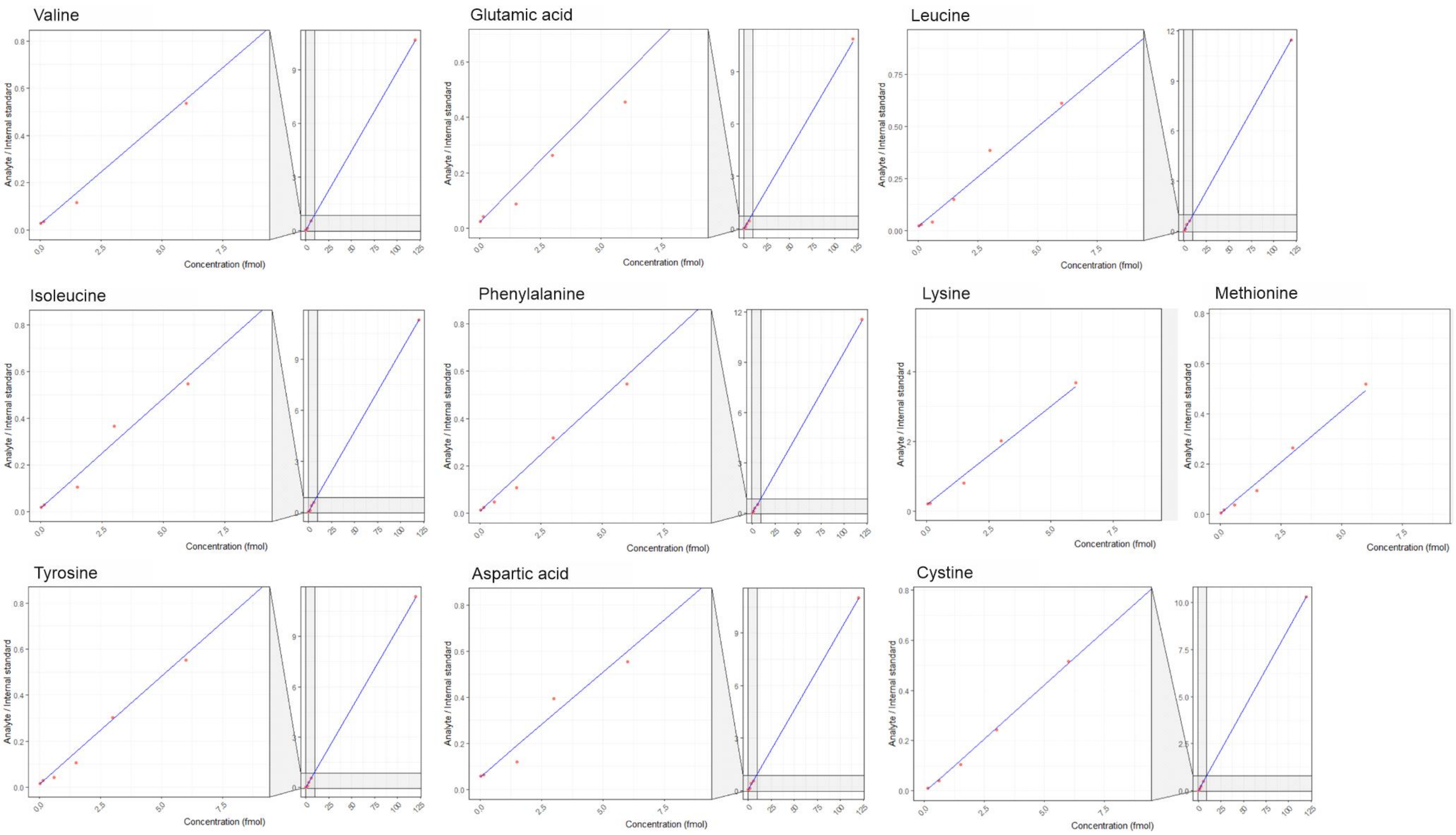

### Figure S2. Retention time comparison on nLC and UHPLC.

Comparison of retention time of nLC and previously[2] performed UHPLC analysis of TMT labeled analytes. The analytes present but unquantifiable (glycine, serine, arginine and histidine) here are also analytes with the lowest retention time in the previous experiments suggesting their poor retention during the sample loading and chromatogram building steps of nLC operation.

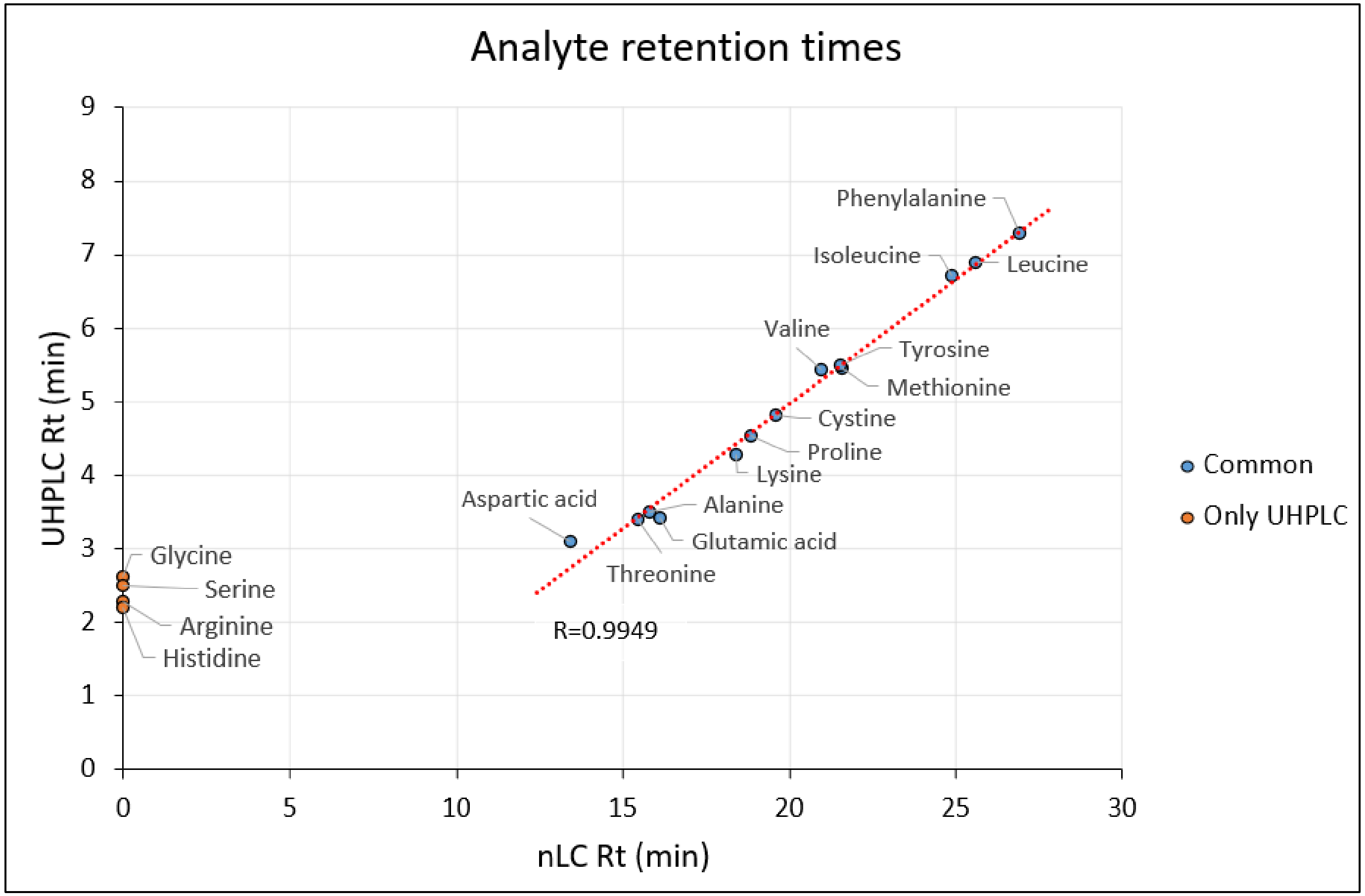

### Figure S3. Heatmaps of single cells.

Heatmap of amine features attained using untargeted metabolomics approach on single cells. Intensities are intra-scan normalized to the carrier. From the 2178 features with cell specific reporter ions along with carrier signal, 249 were common in at least 25% of the samples.

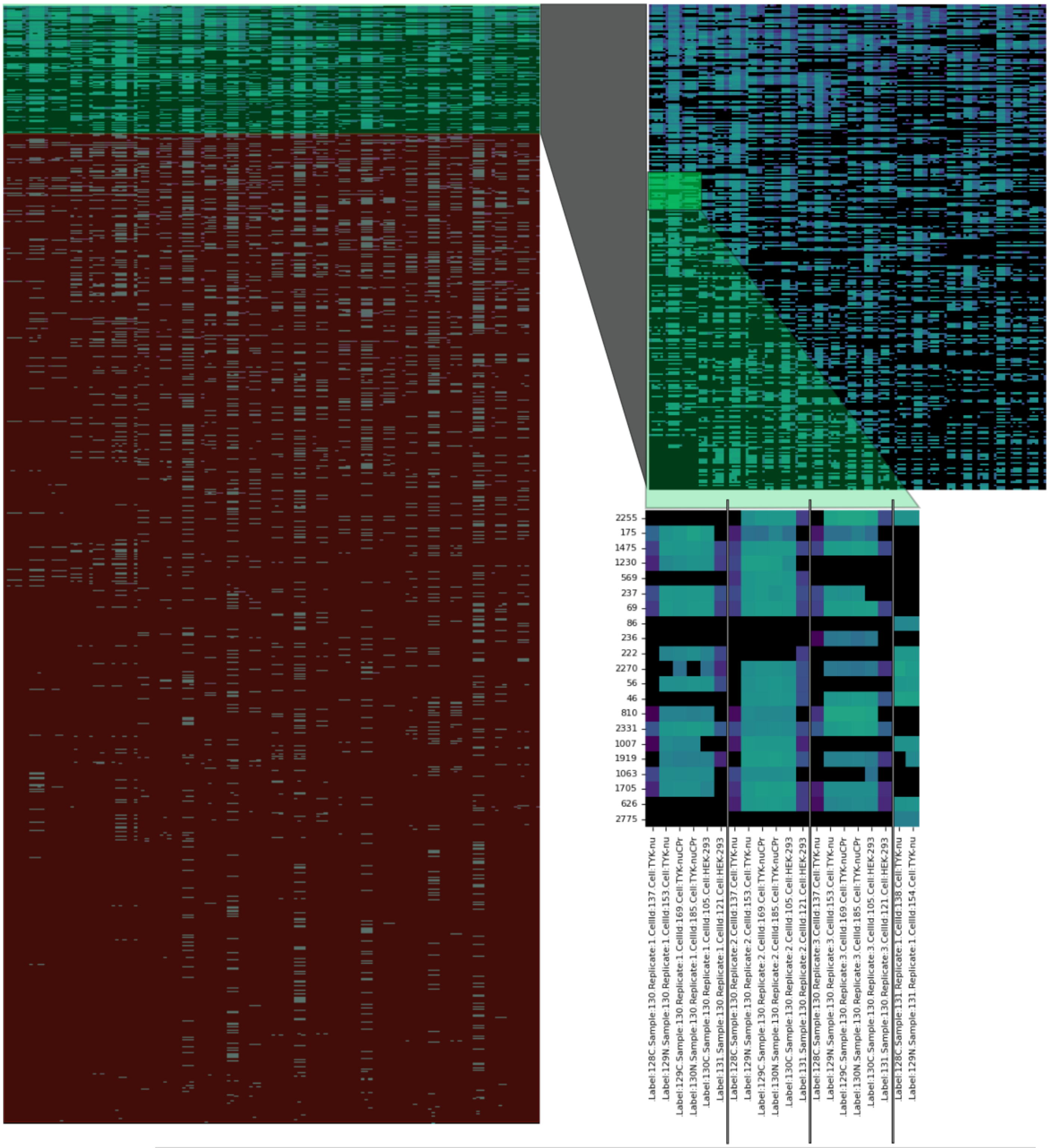

### Figure S4. Example of MS2 scan matching for glutamic acid annotation

Features were annotated using MS2 spectra against previously[2] measured TMT10plex labeled amines (A) and MS2 spectra of unlabeled analyte from GNPS2 database universal spectrum identifier (USI) interface[25] (GNPS Spectrum USI: mzspec_GNPS_GNPS-LIBRARY_accession_CCMSLIB00000081783) with added TMT10plex labeling mass +229.162932 (B). Matches were evaluated using Cosine similarity and proportion of TIC explained by the matching reference ions. To avoid artificially inflating the cosine similarity score with other TMT labeling products only (non-reporter ion) spectra above 132 Da and relative abundance above 0.1% were considered.

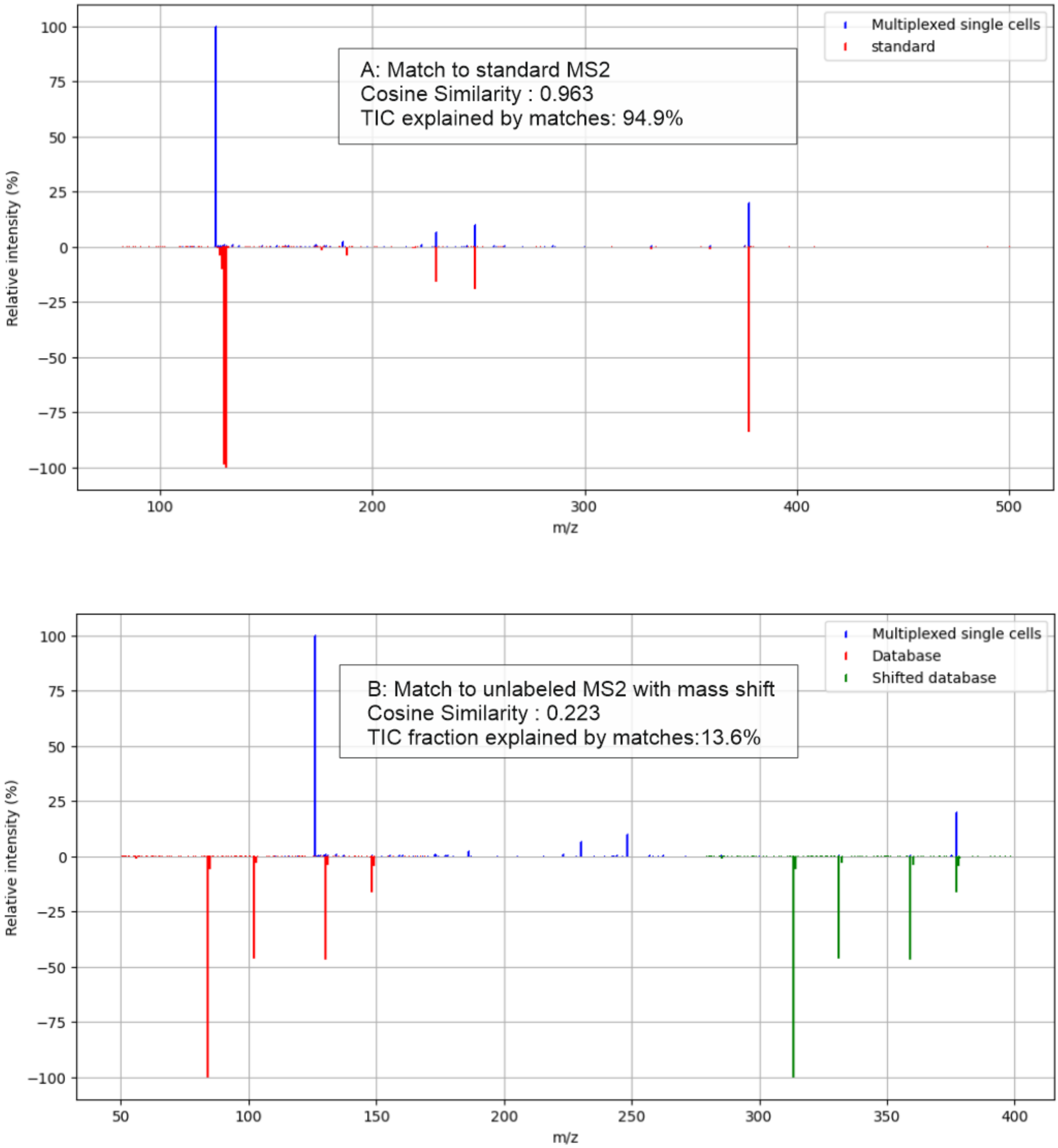

[15] P. Virtanen, R. Gommers, T. E. Oliphant, M. Haberland, T. Reddy, D. Cournapeau, E. Burovski, P. Peterson, W. Weckesser, J. Bright, S. J. van der Walt, M. Brett, J. Wilson, K. J. Millman, N. Mayorov, A. R. J. Nelson, E. Jones, R. Kern, E. Larson, C. J. Carey, İ. Polat, Y. Feng, E. W. Moore, J. VanderPlas, D. Laxalde, J. Perktold, R. Cimrman, I. Henriksen, E. A. Quintero, C. R. Harris, A. M. Archibald, A. H. Ribeiro, F. Pedregosa, P. van Mulbregt, A. Vijaykumar, A. Pietro Bardelli, A. Rothberg, A. Hilboll, A. Kloeckner, A. Scopatz, A. Lee, A. Rokem, C. N. Woods, C. Fulton, C. Masson, C. Häggström, C. Fitzgerald, D. A. Nicholson, D. R. Hagen, D. V. Pasechnik, E. Olivetti, E. Martin, E. Wieser, F. Silva, F. Lenders, F. Wilhelm, G. Young, G. A. Price, G. L. Ingold, G. E. Allen, G. R. Lee, H. Audren, I. Probst, J. P. Dietrich, J. Silterra, J. T. Webber, J. Slavič, J. Nothman, J. Buchner, J. Kulick, J. L. Schönberger, J. V. de Miranda Cardoso, J. Reimer, J. Harrington, J. L. C. Rodríguez, J. Nunez-Iglesias, J. Kuczynski, K. Tritz, M. Thoma, M. Newville, M. Kümmerer, M. Bolingbroke, M. Tartre, M. Pak, N. J. Smith, N. Nowaczyk, N. Shebanov, O. Pavlyk, P. A. Brodtkorb, P. Lee, R. T. McGibbon, R.

Feldbauer, S. Lewis, S. Tygier, S. Sievert, S. Vigna, S. Peterson, S. More, T. Pudlik, T. Oshima, T. J. Pingel, T. P. Robitaille, T. Spura, T. R. Jones, T. Cera, T. Leslie, T. Zito, T. Krauss, U. Upadhyay, Y. O. Halchenko, Y. Vázquez-Baeza, *Nat. Methods* **2020**, *17*, 261–272.

[23] M. Wang, J. J. Carver, V. V. Phelan, L. M. Sanchez, N. Garg, Y. Peng, D. D. Nguyen, J. Watrous, C. A. Kapono, T. Luzzatto-Knaan, C. Porto, A. Bouslimani, A. V. Melnik, M. J. Meehan, W. T. Liu, M. Crüsemann, P. D. Boudreau, E. Esquenazi, M. Sandoval-Calderón, R. D. Kersten, L. A. Pace, R. A. Quinn, K. R. Duncan, C. C. Hsu, D. J. Floros, R. G. Gavilan, K. Kleigrewe, T. Northen, R. J. Dutton, D. Parrot, E. E. Carlson, B. Aigle, C. F. Michelsen, L. Jelsbak, C. Sohlenkamp, P. Pevzner, A. Edlund, J. McLean, J. Piel, B. T. Murphy, L. Gerwick, C. C. Liaw, Y. L. Yang, H. U. Humpf, M. Maansson, R. A. Keyzers, A. C. Sims, A. R. Johnson, A. M. Sidebottom, B. E. Sedio, A. Klitgaard, C. B. Larson, C. A. P. Boya, D. Torres-Mendoza, D. J. Gonzalez, D. B. Silva, L. M. Marques, D. P. Demarque, E. Pociute, E. C. O'Neill, E. Briand, E. J. N. Helfrich, E. A. Granatosky, E. Glukhov, F. Ryffel, H. Houson, H. Mohimani, J. J. Kharbush, Y. Zeng, J. A. Vorholt, K. L. Kurita, P. Charusanti, K. L. McPhail, K. F. Nielsen, L. Vuong, M. Elfeki, M. F. Traxler, N. Engene, N. Koyama, O. B. Vining, R. Baric, R. R. Silva, S. J. Mascuch, S. Tomasi, S. Jenkins, V. Macherla, T. Hoffman, V. Agarwal, P. G. Williams, J. Dai, R. Neupane, J. Gurr, A. M. C. Rodríguez, A. Lamsa, C. Zhang, K. Dorrestein, B. M. Duggan, J. Almaliti, P. M. Allard, P. Phapale, L. F. Nothias, T. Alexandrov, M. Litaudon, J. L. Wolfender, J. E. Kyle, T. O. Metz, T. Peryea, D. T. Nguyen, D. VanLeer, P. Shinn, A. Jadhav, R. Müller, K. M. Waters, W. Shi, X. Liu, L. Zhang, R. Knight, P. R. Jensen, B. Palsson, K. Pogliano, R. G. Linington, M. Gutiérrez, N. P. Lopes, W. H. Gerwick, B. S. Moore, P. C. Dorrestein, N. Bandeira, *Nat. Biotechnol.* **2016**, *34*, 828–837.

[24] E. L. Schymanski, J. Jeon, R. Gulde, K. Fenner, M. Ruff, H. P. Singer, J. Hollender, *Environ. Sci. Technol.* **2014**, *48*, 2097–2098.

[25] W. Bittremieux, C. Chen, P. C. Dorrestein, E. L. Schymanski, T. Schulze, S. Neumann, R. Meier, S. Rogers, M. Wang, *bioRxiv* **2020**, 2020.05.09.086066.
